# Earlier flowering explains only a small part of experimental drought’s effects on a wildflower’s population growth

**DOI:** 10.64898/2026.03.25.714308

**Authors:** Scott W. Nordstrom, Jenna A. Loesberg, Piper Battersby, Jennifer L. Williams

**Affiliations:** Department of Geography, University of British Columbia, Vancouver, BC, V6T1Z2; Biodiversity Research Centre, University of British Columbia, Vancouver, BC, V6T1Z4; Population Research Center, Portland State University, Portland, OR, 97201

**Keywords:** integral projection model, rainfall manipulation, phenology

## Abstract

Flowering times are shifting with climate change, but whether these changes will affect population dynamics of long-lived plants remains largely unknown. We combined eight years of demographic censuses and a four-year phenological survey to examine the effects of experimental drought and irrigation on flowering phenology and population growth in a perennial wildflower using integral projection models. Drought advanced flowering and earlier-flowering plants produced more seeds regardless of treatment. However, when decomposing rainfall treatment effects on population growth rates into effects of phenology and direct, non-phenological effects, direct effects on both growth and reproduction were much larger than phenology-driven changes in seed production for either treatment. Our results highlight how phenological shifts may only marginally impact population dynamics in perennial plants and demonstrate that assessing flowering phenology’s consequences for persistence under climate change must also account for direct demographic effects of the climate driver itself.

## Introduction

The timing of life cycle events in plants and animals is shifting with climate change, providing an important and well-documented indicator of climate change’s effects (Parmesan 2007, Iler et al. 2021, Inouye 2022). Plants flowering earlier in response to warming is one of the most prominent phenological shifts (CaraDonna et al. 2014, Collins et al. 2021, Collins et al. 2025). Flowering time responses to precipitation change are more varied, including earlier (Kooyers 2015, König et al. 2018, Van Dyke & Kraft 2025), later (Moore & Lauenroth 2017, Castillioni et al. 2021), or no change in response to drought (Reed et al. 2019, Kuppler & Kotowska 2021, Brunet et al. 2025). Changing phenology can influence demographic processes such as seed production (reviewed in Iler et al. 2021). However, for longer-lived plants, whether these shifting flowering times affect population growth, and thus have the potential to affect population persistence, is not well understood (Iler et al. 2021, Zettlemoyer & DeMarche 2022). Considering the reality of rapid and continuing climate change, understanding the role of phenology in population viability is critical for ecology and conservation.

Understanding how phenology contributes to population viability requires not only linking phenology to individual demographic processes, but also quantifying population growth rates and their sensitivities to flowering times (e.g., Ehrlén & Münzbergova 2009, Knight et al. 2009). Few studies have done this in the context of climate change (Iler et al. 2021). Those that have primarily studied montane or alpine populations and focused exclusively on phenological effects of warming or snowmelt timing (Campbell 2019, Iler et al. 2019, Keller & Shea 2021). This leaves other ecosystems and potential drivers of phenology such as drought (Kooyers 2015, Kazan & Lyons 2016) critically understudied. Furthermore, existing studies of phenology and population growth feature at most one year of experimental manipulation (Iler et al. 2019, Keller & Shea 2021). When a vital rate (e.g., seed production per plant) changes, the expected change to the population growth rate is proportional to the magnitude of the vital rate change and the sensitivity of the population growth rate to the vital rate (Caswell 1989). For long-lived plants, prior demographic and life history results suggest that flowering phenology’s influence on population dynamics could be small. Population growth in long-lived plants tends to be less sensitive to changes in reproduction than to changes in other vital rates, e.g., survival or growth (Franco & Silvertown 2004), suggesting that substantial changes in seed production are needed for reproductive phenology to influence population growth. Additionally, other vital rates may change in concert with or due to shifting flowering phenology (Ehrlén 2015); if these changes occur in opposing directions, their effects may partially or entirely offset each other, leaving population growth rates relatively stable as the climate changes (Villellas et al. 2015).

At the same time that climate may be affecting fitness through phenological shifts, it may be directly altering other demographic processes, e.g., through physiological stress (Wang & Xu 2026). For example, phenology-driven changes to seed production may be compounded or counteracted by climate-driven changes to flower size or number (Kuppler & Kotowska 2021, Brunet et al. 2025). Analysis of climate-mediated phenological shifts on fitness must account for these potentially confounding effects (Fig. 1). Recent work has considered or explicitly quantified distinct contributions of warming on seed production either directly or mediated by earlier flowering (Zettlemoyer et al. 2024, Zhang et al. 2024, Collins et al. 2025), finding varied results. To our knowledge, this distinction has not yet been applied to population dynamics or lifetime fitness. This oversight potentially obscures the true mechanisms underlying the effects of climate and phenology on population persistence.

**Figure 1.**
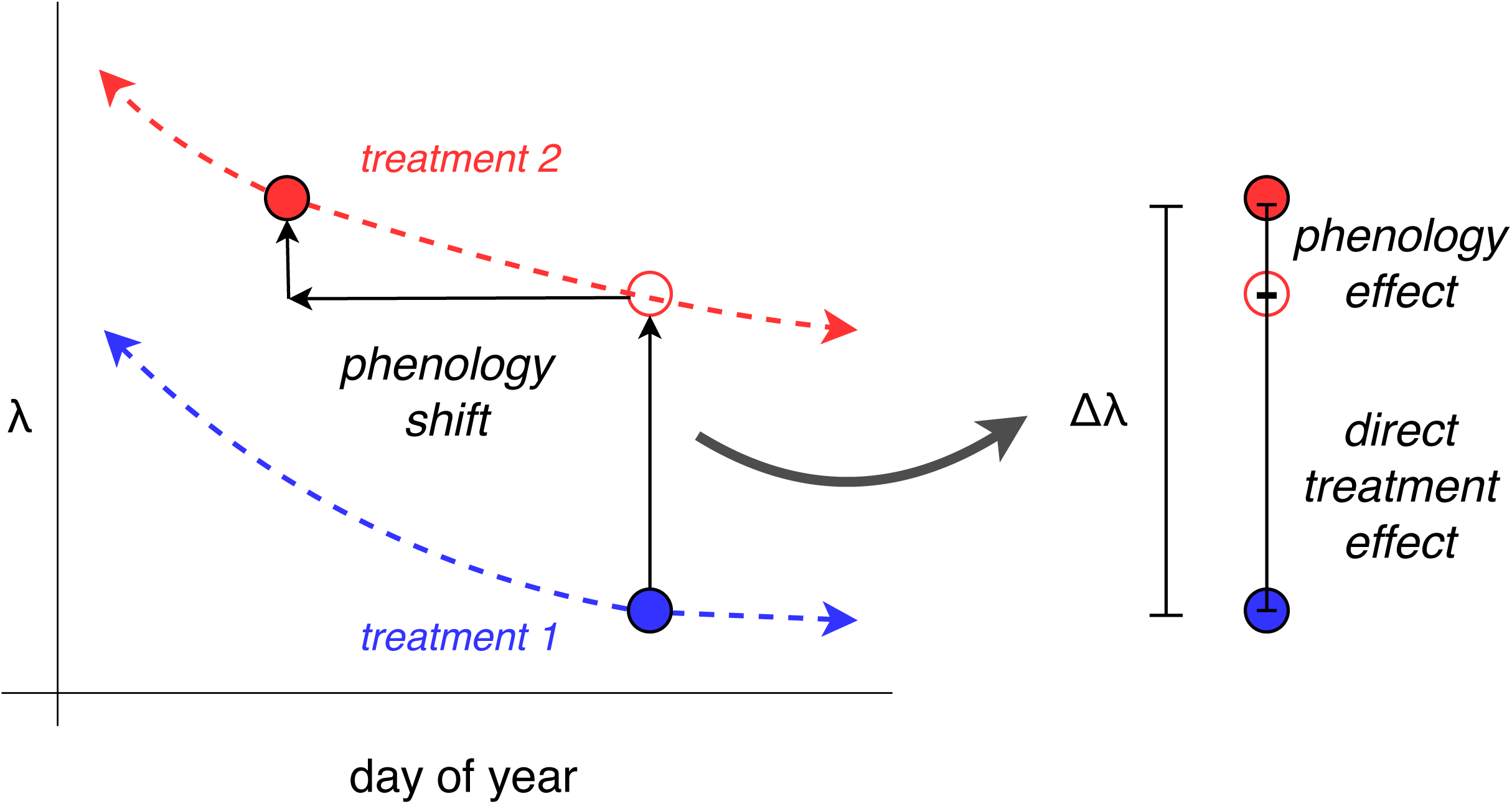
Population growth rates (λ) can be affected independently by phenology and climate drivers. Here, solid circles represent estimates of λ for two hypothetical climate treatments on their mean flowering dates; dashed curves estimate λ on other flowering dates. Vertical arrows are estimated treatment and phenological effects, respectively, of treatment 2 compared to treatment 1. See Supporting Information for technical details.

We quantified the effects of changes in rainfall and flowering phenology on population growth in a common perennial herb, *Lomatium utriculatum*. We established an *in-situ* rainfall manipulation experiment in oak savanna on Vancouver Island, BC, Canada, to explore the effects of drought and increased rainfall (*via* irrigation) on demography; both treatments are relevant to this ecosystem, as projections predict both increased annual precipitation and increasingly dry summers (Cowichan Valley Regional District 2017). We present results from eight years of demographic censuses and four years of phenological surveys. These observations from our multi-year manipulative experiment allow us to infer causality and robustly quantify responses of flowering phenology and vital rates to changes in rainfall in a natural setting (Miller-Rushing et al. 2010, Williams et al. 2025). We ask the following questions: (1) do changes in rainfall influence the flowering phenology of *L. utriculatum*, (2) does changing rainfall influence vital rates, either through flowering phenology or directly, and (3) how does changing rainfall influence population growth, either through flowering phenology or directly (*sensu* Fig. 1)? We answer this final question using integral projection models (Merow et al. 2014) and a two-way life table response experiment (Caswell 1989).

## Materials and methods

### Study system

*Lomatium utriculatum* (Apiaceae) is a tap-rooted perennial herb that grows in grassland and savanna habitat ranging from southern California to southern British Columbia. We studied a population within a Garry oak savanna preserve on Vancouver Island near Duncan, BC (48.809, -123.631); this region has a Mediterranean climate where cool, wet winters and springs are followed by warm, dry summers. *Lomatium utriculatum* is locally common in these savannas, which are characterized by Garry Oak (*Quercus garyanna*) trees with a diverse understory of grasses and forbs. Prior to European settlement, savannas were managed for millennia by First Nations *via* prescribed burning (McCune et al. 2013).

*Lomatium utriculatum* reproduces sexually with discrete inflorescences (umbels). Flowering is indeterminate within growing seasons, which typically begin in March and are constrained by the onset of the dry season beginning in late June at this site. Individual inflorescences are frequently aborted or grazed. We established a rainfall manipulation experiment in 2015 following the International Drought Experiment protocol (Knapp et al. 2017, Smith et al. 2024), with five permanent 3x3 m^2^ drought shelters that reduce incipient precipitation by 50% to simulate a one-in-100-year drought (Smith & Williams, 2023). Fifteen 2x1 m plots were established to collect demographic data, with five beneath drought shelters, five passively irrigated with collected rainfall during the growing season, and five unmanipulated control plots (Smith & Williams 2023). During 2021-2024, we added an additional 15–22.5 mm/m^2^ of water per week during the spring growing season due to previously dry springs (Loesberg & Williams 2025). We observed *L. utriculatum* in 14 plots during the study.

### Data collection and processing

We collected demographic data annually starting in 2016 (when each *L. utriculatum* individual was assigned a permanent metal tag), assessing survival and recording number of leaves, length of longest leaf, and number of inflorescences. New individuals received tags, and if an individual was not observed in two successive years it was presumed dead. We imputed survival for 484 records (11% of records of surviving plants 2017-2023) where plants were not observed but observed alive the following year, assuming the plant was missed due to observer error, herbivory, or vegetative dormancy.

*Lomatium utriculatum* produces inflorescences in the form of compound umbels with small, yellow-petaled flowers. We defined an inflorescence’s flowering date as the date we first observed yellow petal tissue visible (Fig. 2a). We chose this definition because tissue visibility was the least ambiguous measure of flowering, increasing consistency among years. In 2021-2024, we conducted weekly flowering phenology surveys recording the number of budding or developed inflorescences with yellow floral tissue visible. All flowering plants were included, except in 2023 when a subset was excluded due to large numbers of flowering plants in some plots. Because we did not uniquely label individual inflorescences, we estimated phenology at the plant level using the mean flowering date across inflorescences on the plant. Prior to seed dispersal (early-mid June), we counted the number of seeds per inflorescence, except in 2021 when seed was counted for 1-2 randomly-sampled inflorescences per plant. Although each plot was censused once per week, they were not censused on the same day each week; 88% of plant resurvey dates followed the most recent visit by 5-9 days, and only 10% of the 438 plant resurveys occurring fewer than five or more than nine days after their previous survey included a new inflorescence appearing on the plant. To mitigate potential bias, all survey dates within a year were coded to the first day of each week such that each survey date was seven days apart. While this mitigates potential bias within years, it does not mitigate (or may even introduce) spurious effects among years; for this reason, we do not interpret inter-annual variation in phenology estimated in our models.

**Figure 2:**
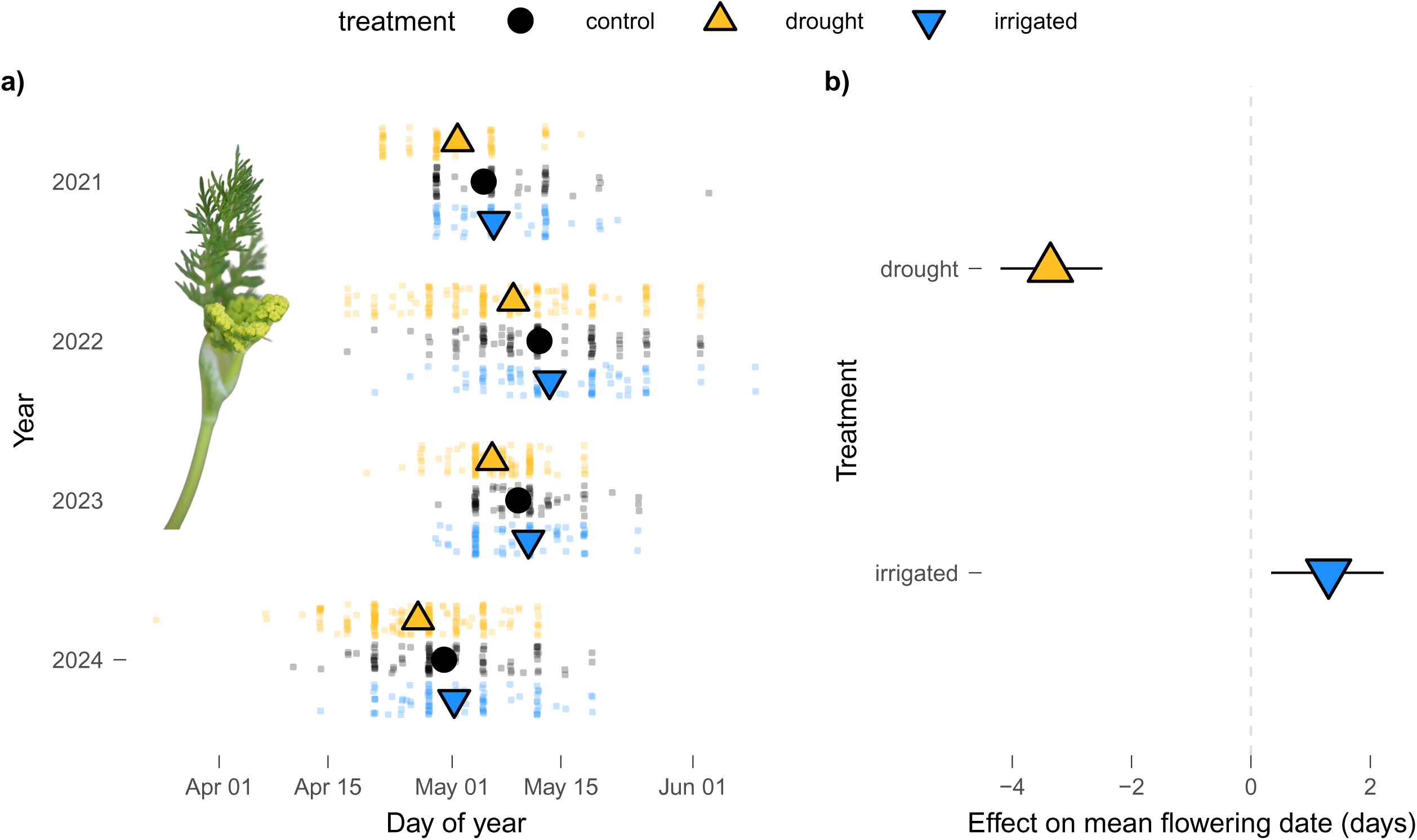
Rainfall manipulation effects on flowering phenology. a) Plant mean flowering dates for each plant over the course of the study, with example of an emerging inflorescence with yellow tissue visible in left inset. Small points each correspond to one plant, jittered by half a day to increase visibility. Large points are treatment means in each year as estimated by our best-supported model. b) Mean effect of each rainfall manipulation treatment on plant flowering date, relative to control treatment. Segments are 95% confidence intervals estimated with 500 bootstrapped samples.

### Phenology models

We evaluated the effects of rainfall manipulation on phenology and vital rates by fitting candidate mixed-effects models with and without categorical treatment effects and selecting the most parsimonious model with ΔAIC<2 of the lowest-AIC model. Models estimating treatment effects on phenology used the mean Julian flowering date of plants in each year as a continuous response with Gaussian errors. The best-supported phenology model included treatment but no interaction with year (ΔAIC=0.98, Table S2). We used this model to estimate a mean flowering date for each treatment (hereafter, *x̅*_*j*_ for treatment *j*), obtaining a single date per treatment by averaging predicted annual means across the four years of study. While rainfall treatment shifted phenology on average, the large degree of individual variation meant that treatment explained only a small (<10%) portion of variance in flowering time (SI Results); this phenological variation independent of rainfall manipulation allowed us to separate treatment from phenology effects in vital rate models. Treatment and vital rate relationships with phenology were qualitatively identical when modeling phenology as a plant’s first flowering day rather than its mean (Tables S4-5).

### Vital rate models

We modeled the *L. utriculatum* life cycle with eight vital rates: survival, individual growth (conditioned on survival), flowering (binary outcome), number of inflorescences (conditioned on flowering), probability of inflorescence surviving to seed (henceforth, inflorescence “success”), seeds per successful inflorescence, seedling establishment probability, and size of new recruits (Table S1). All vital rates were estimated as continuous functions of plant size (log_e_(length of longest leaf x number of leaves)), with two exceptions: recruit size, where size was the response, and seedling establishment probability, where we chose a value (∼0.0046) that fixed λ=1 for the control treatment. This establishment probability was within a 95% confidence interval of an estimate from a one-year seed addition study (SI Methods). Our results were qualitatively similar with other establishment probabilities values (Figs. S4-5), with declining precision with higher establishment rates. Survival was modeled with a logistic generalized linear mixed effects model (GLMM), growth was modeled with a linear mixed effects model (LMM), probability of flowering and number of umbels per flowering plant were jointly estimated using a zero-truncated Poisson GLMM, umbel success and seed count were jointly estimated using a zero-inflated negative binomial GLMM, and recruit size was modeled with an LMM.

For all vital rates aside from seed germination, we evaluated whether rainfall treatment improved candidate models with size and year present in the model. Using phenology and seed counts from 2021-2024, we also tested for effects of plant mean flowering date on inflorescence success and seed count, and for effects of phenology on growth and survival of plants in the subsequent year. We fit candidate models with linear and quadratic effects of phenology and treatment-phenology interactions. See SI Methods and SI Results for more detail, including model selection and final model coefficients.

### Integral projection models

Best-supported vital rate models were combined to fit integral projection models (Merow et al. 2014), defined by

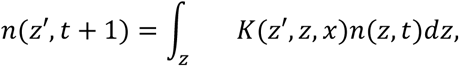

where *K(z’,z,x)* is the inter-annual transition kernel from size *z* to *z’* for a population with mean flowering date *x* (shifted such that *x=*0 mean flowering date observed in control plots), and *n(z,t)* gives the density of individuals between size *z* and *z+*Δ*z* in time step *t*. The kernel is a product of survival-growth and reproductive subkernels, i.e., *K(z’,z,x)=T(z’,z,x)R(z’,z,x)*. The survival-growth subkernel is defined as *T(z’,z,x)=s(z)g(z’,z,x)* for survival probability *s(z)* (found in model selection to be independent of prior year’s flowering date) and growth function *g(z’,z,x)*. The growth function is a normally distributed probability density function (PDF) with mean

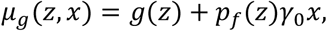

where *g(z)* is the expected size at time *t* of a surviving individual of size *z* at *t-1*, *p_f_(z)* is the probability of flowering, *γ_0_* is the per-day effect of prior-year phenology on growth, and *x* the size of the phenological shift (relative to the mean flowering date in control plots). The standard deviation of the PDF is the Gaussian residual term of the growth vital rate regression.

The reproductive kernel is defined by

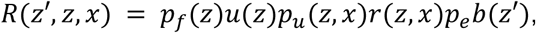

where *p_f_(z)* is the probability of flowering, *u(z)* is the number of inflorescences produced per flowering plant, *p_u_(z,x)* is the per-inflorescence success probability for a plant with mean flowering date *x*, *r(z,x)* is the number of seeds produced per successful inflorescence, *p_e_* is the probability of seedling establishment, and *b(z’)* is a normal PDF for the size of new recruits. For a given flowering date *x* and treatment *j*, we estimated the kernel *K(z’,z*,*x*) with all vital rates evaluated at the given *x* and *j*, and estimated the asymptotic deterministic growth rate, *λ(x,j)*, by taking the dominant eigenvalue of this kernel.

We used these kernels for two sets of estimates. First, we estimated the sensitivity of λ to flowering date within each treatment by estimating λ for each treatment at flowering dates across a four-week period centered at the mean flowering date in the controls (i.e., with *x* ranging from -14 to 14). Second, we estimated a single λ for each treatment by evaluating kernels at the mean flowering date of that treatment, i.e., by estimating *λ*(*x̅*_*j*_, *j*) for each *j.* These *λ*(*x̅*_*j*_, *j*) are used in the LTRE analysis, described below.

### Life Table Response Experiment

To determine the relative contributions of shifts in flowering time and rainfall treatment on λ, we used a life table response experiment (LTRE; Caswell 1989). Specifically, we used a two-way LTRE to decompose Δλ between rainfall treatments into contributions of phenological shifts (*x*) and direct treatment effects (*j; sensu* Fig. 1) acting through each vital rate. This analysis is retrospective and analyzes observed differences in λ between treatments rather than projecting future dynamics (Caswell 2000). Our LTRE estimates contributions from each individual vital rate parameter that varies between treatments to the overall treatment effect on λ. The magnitude of a vital rate parameter’s contribution is the product of the parameter difference between treatments and the sensitivity of λ to that parameter (Caswell 1989, Griffith 2017). We define *φ^(j)^* and *ψ^(j)^* as the respective contributions of phenology- and direct treatment-driven changes to vital rate *Y* to the effect of treatment *j* on λ (see Supporting Information for formal definition and derivation). Summing *φ* and *ψ* terms over all vital rates approximates Δλ, i.e., for treatment *j*,

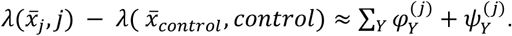

Our LTRE is a first-order decomposition; we elected not to estimate higher-order effects (interactions between phenology and treatment) for simplicity and because our first order LTRE had accuracy within 3% of the true Δλ (see Supporting Information). An alternative formulation of our LTRE with order of operations reversed (see Supporting Information) produced qualitatively identical results (Table S19). We estimated sensitivity of λ to each vital rate parameter by perturbing the parameter and measuring the change in λ in the resulting kernel (approach 5 in Griffith 2017). For vital rates models where both slope and intercept varied by treatment, the treatment effect was the sum of slope and intercept contributions. Vital rate parameters that do not vary with phenology or treatment have contributions equal to zero and are not included in results.

### Estimate uncertainty

We estimated uncertainty in rainfall treatment effect on phenology, phenology or treatment effects on vital rates, λ, and LTRE components using bootstrapped resampling. Specifically, we re-estimated the model of treatment effects on phenology and each vital rate model with a treatment or phenology effect with 500 datasets re-sampled with replacement. We then used this ensemble of 500 bootstrapped phenology and vital rate estimates to re-fit kernels, used both to provide uncertainty in λ and to fit 500 bootstrapped LTREs, capturing uncertainty in LTRE contributions. Bootstrapped *p*-values are defined as the fraction of bootstrapped estimates with the opposite sign of the point estimate; all reported *p-*values are bootstrapped values unless otherwise noted. All analysis was performed in R v.4.5.2 (R Core Team 2023) with models fit using *glmmTMB* v1.1.13 (Brooks et al. 2017).

## Results

### Rainfall treatment effects on phenology

Rainfall manipulation shifted mean plant flowering dates (Fig. 2; Likelihood-Ratio test for model with vs. without treatment p=0.004). Under drought, flowering occurred 3.4 days earlier on average (bootstrapped p<0.002), while increased rainfall (irrigation) delayed flowering by 1.3 days (p=0.006; Fig. 2b). Across four years, mean annual flowering dates spanned two weeks (April 29 to May 10). Treatment effects accounted for roughly 7% of unexplained variance after accounting for inter-annual variation (pseudo-R^2^ of model without treatment: 0.27; with treatment: 0.32; Nakagawa & Schielzeth 2013; see our Supporting Information).

### Phenology and treatment effects on vital rates

Inflorescence success and seeds per successful inflorescence varied with a plant’s mean flowering date, but in opposing directions (Fig. 3). Earlier flowering increased inflorescence mortality (e.g., due to abortion; Fig. 3a) while successful inflorescences on earlier-flowering plants produced more seeds (Fig. 3b). The net effect was that earlier-flowering plants produced more seeds (Fig. 3c): an average-sized flowering plant in a control plot flowering one week earlier produced ∼2 more seeds per inflorescence (5.5% increase). Phenological effects did not vary by rainfall treatment for either reproductive component (both L-R tests for treatment-phenology interactions: p>0.4). Likewise, there was no strong evidence for quadratic effects of flowering date on either reproductive component, suggesting that neither component has an intermediate optimal flowering time (L-R test for quadratic phenology term on umbel success: p=0.89; for quadratic term on seeds per successful umbel: p=0.07). Plants flowering early in the prior year grew more than their later-flowering counterparts (Fig. 3d). Survival did not vary by flowering date in the prior year.

**Figure 3.**
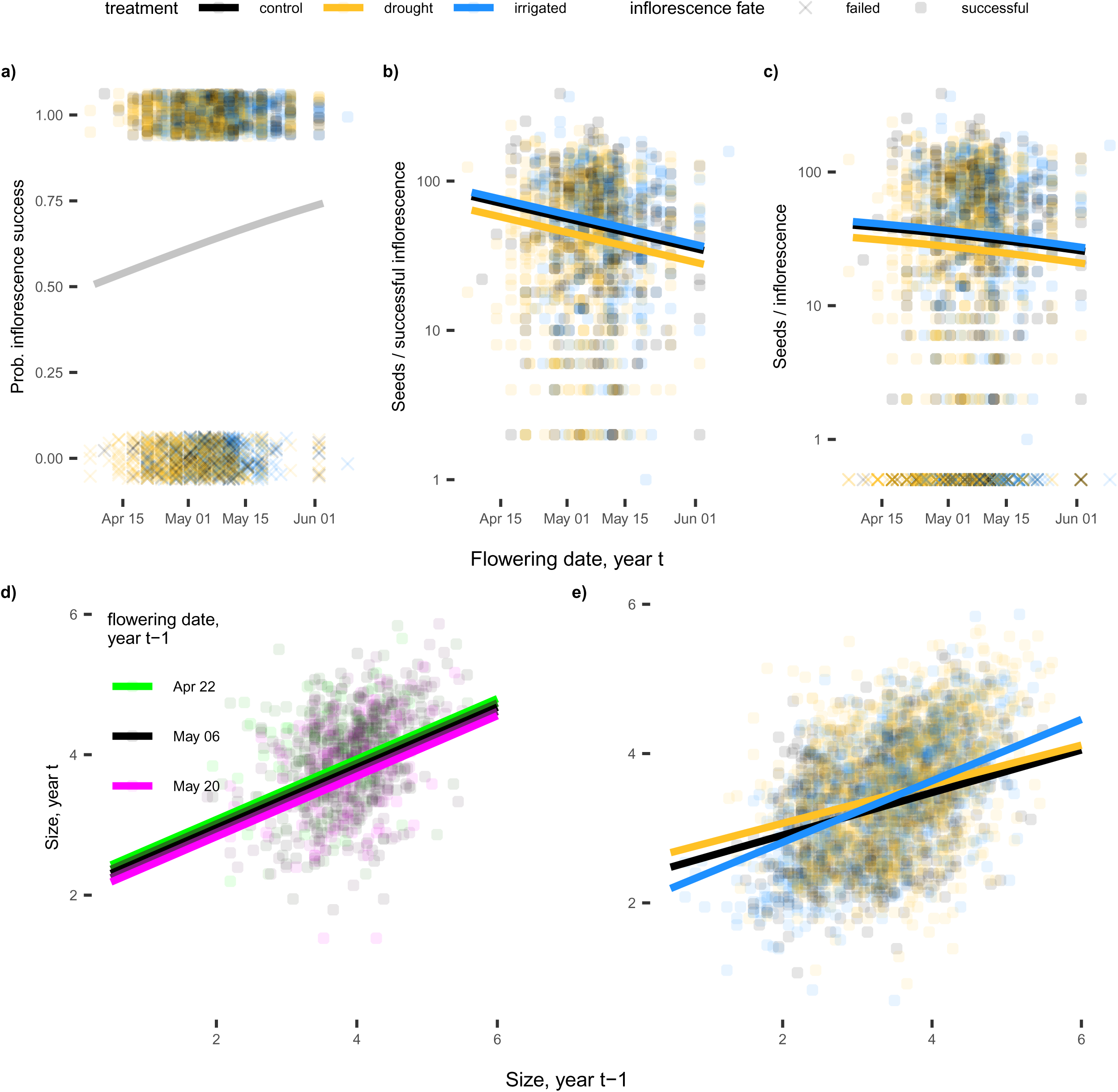
Phenology-mediated and direct rainfall treatment effects on components of reproduction (a-c) and growth (d-e). a) Inflorescence success by flowering date. Grey curve denotes no differences among treatments. b) Seeds per successful inflorescence by flowering date. c) Total seeds per inflorescence by flowering date (those producing zero seeds along bottom). Curves in a-c are for a plant with mean size of flowering plants in 2021-2024 and in c) are the products of curves in a) and b). d) Individual plant growth, conditioned on flowering date in prior year. e) Individual plant growth by treatment. One point represents one inflorescence in panels a-c and one plant in panels d-e.

Both rainfall treatments reduced overall seed count even after statistically controlling for the effects of plant phenology (Fig. S3), but through different demographic pathways. Drought reduced the number of seeds produced per inflorescence (Figs. 3b, S3c). Irrigation increased the number of seeds per inflorescence for small plants (Fig. S3c) but reduced the probability of flowering for all sizes (Fig. S3a). In contrast to reproduction, both rainfall manipulation treatments enhanced individual plant growth. Drought increased growth for medium-sized and large plants while irrigation reduced growth of small plants and increased the growth of large plants (relative to growth in controls; Fig. 3e, S2, S3). New recruits were ∼10% larger under drought than plants in control plots. Rainfall treatment did not directly affect survival, number of inflorescences per plant, or probability of umbel success (Fig. S3).

### Phenology and treatment effects on λ

Earlier flowering increased the deterministic population growth rate (λ); flowering one week earlier than the mean flowering date during the study (May 6) increased λ by 0.010 for each treatment (Fig. 4a). Although estimates of λ were higher in both treatments compared to controls, these differences were not significant (Δλ=0.015, bootstrapped p=0.098, Δλ=0.005, p=0.204, for drought and irrigation, respectively). Extrapolating across a four-week period, λ was consistently higher in drought and irrigation treatments compared to controls, although these differences were not statistically significant for either treatment (Fig. 4b).

**Figure 4.**
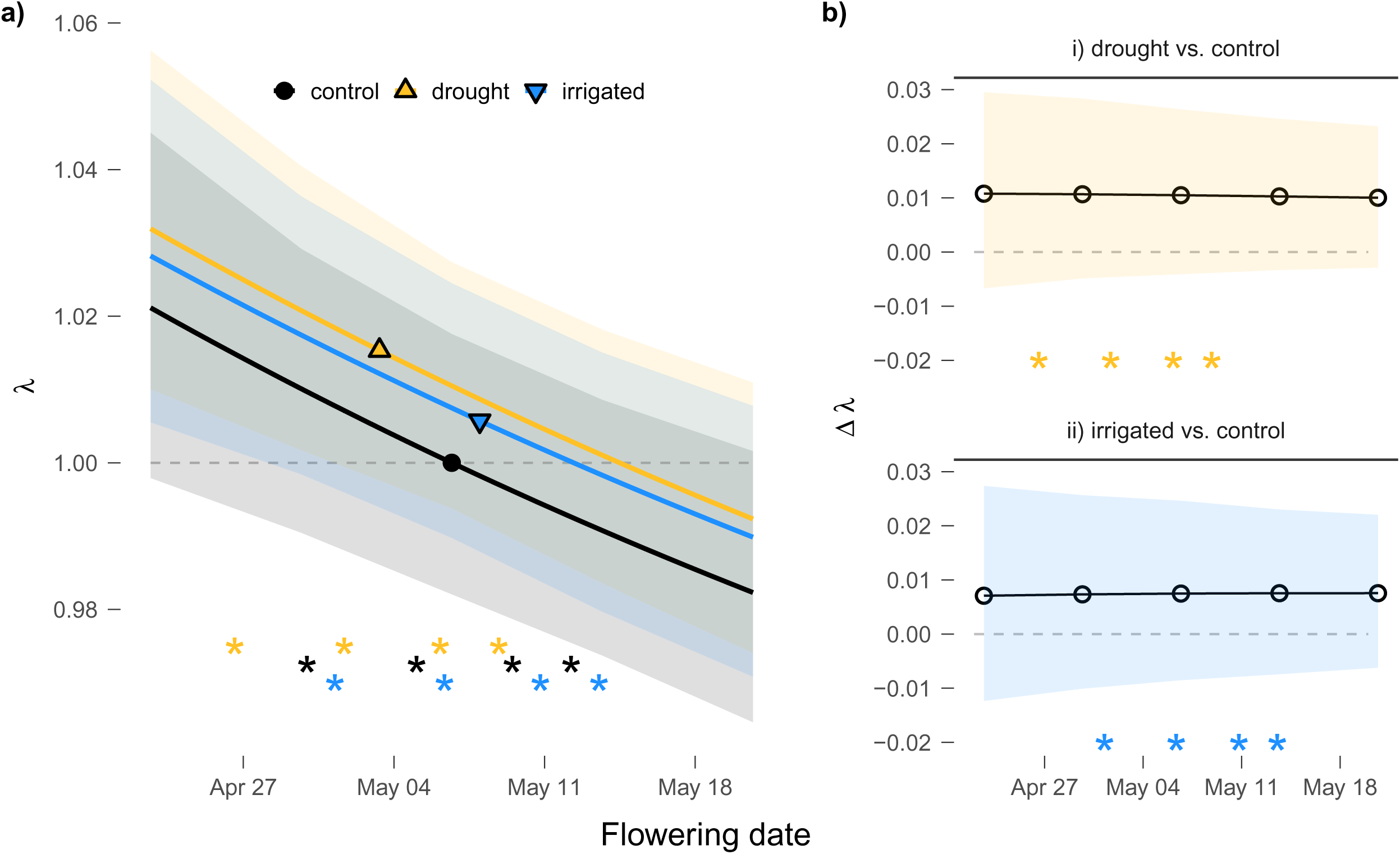
The responsiveness of population growth rates (λ) to flowering date and experimental rainfall over the flowering season. a) λ estimated over a four-week period. Solid shapes give λ for each treatment on the treatment’s mean flowering date. Shaded areas are 95% bootstrapped confidence intervals. b) Differences in λ (Δλ) between control and i) drought and ii) irrigation on the same flowering date, estimated at weekly intervals over the same period. Open circles are mean estimates and shaded regions are 95% bootstrapped confidence intervals. In both panels, asterisks show annual mean flowering dates per treatment during 2021-2024.

Under both rainfall treatments, λ increased primarily due to increased individual growth (Fig. 5). Earlier flowering under drought increased λ through increased seed production, but this effect was small and negated by the drought treatment’s larger direct reduction in seed (*ψ_seed_*, Fig. 5a). For both rainfall treatments, phenology’s combined contribution to Δλ through reproduction (*φ_succ_* + *φ_seed_*) was smaller than phenology’s effects on growth (*φ_grow_*) and direct treatment effects on reproduction (*ψ_succ_* + *ψ_seed_*, Fig. 5b). Survival did not vary with phenology or treatment and thus did not contribute to Δλ.

**Figure 5.**
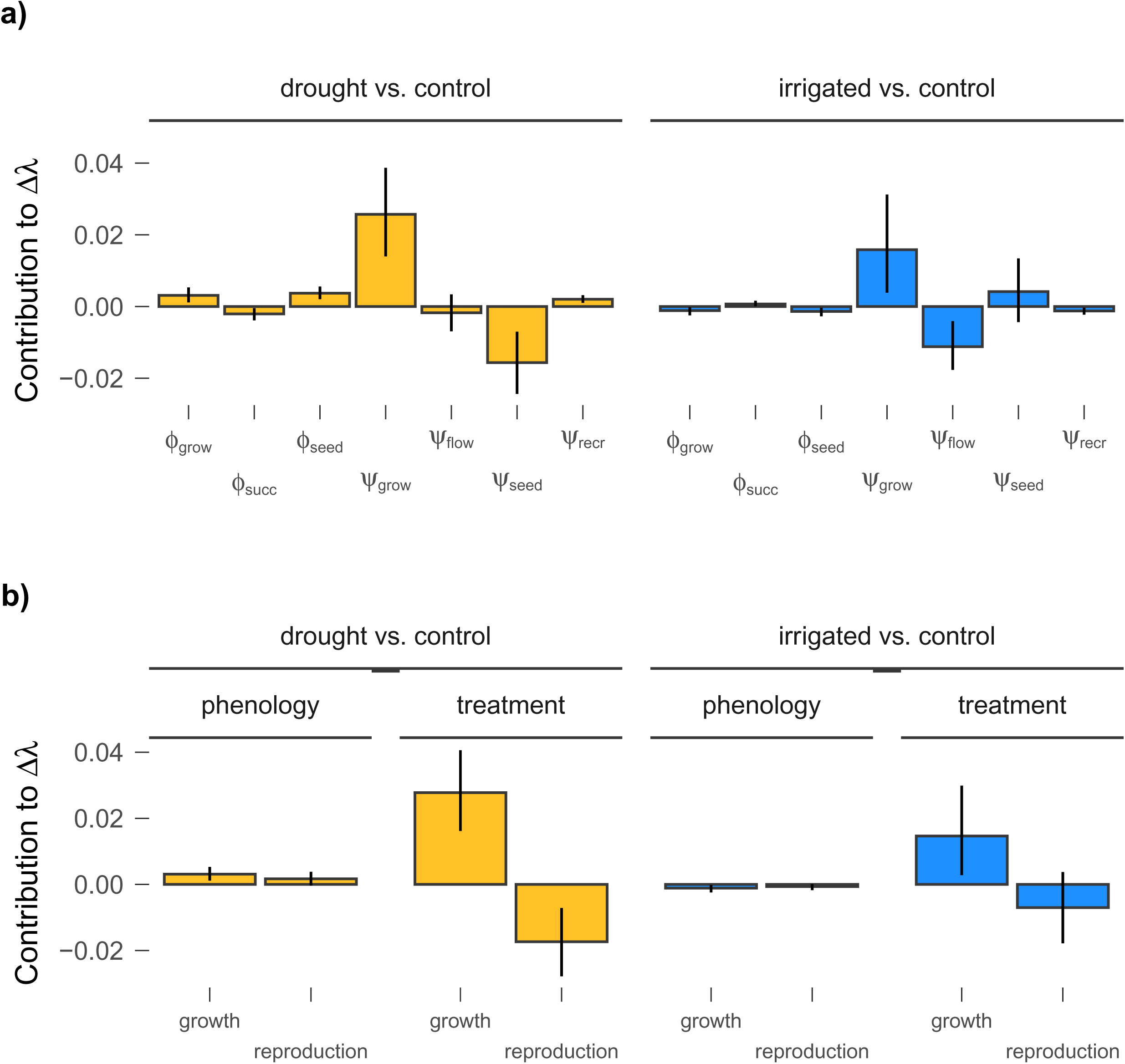
Contributions to differences in λ from phenology-driven and direct treatment effects estimated by a two-way LTRE. a) Contributions from individual vital rates. φ denotes contributions to Δλ through changes in the vital rate through shifting flowering times; ψ denotes contributions to Δλ through non-phenological direct treatment effects on the vital rate. Subscripts: *grow* is individual growth, *succ* is inflorescence success, *seed* is seed production, *flow* is probability of flowering, *recr* is size of new recruits. b) Contributions to Δλ aggregated into effects acting through phenology or treatment (i.e., summing φ or ψ terms, respectively) and changes to growth-related vital rates (individual growth and recruit size) or reproductive vital rates (flowering probability, inflorescence success, and seed count). In both panels, vertical lines are 95% bootstrapped confidence intervals.

## Discussion

We quantified sensitivity of population growth of the common perennial wildflower *Lomatium utriculatum* to experimental rainfall manipulation, considering both changes attributable to changing phenology and those directly influenced by drought or increased rainfall. Reduced rainfall advanced average flowering times by several days, and plants that flowered earlier produced slightly more seeds on average. However, regardless of their flowering date, plants under both rainfall reduction or addition produced fewer seeds compared to controls. Additionally, and surprisingly, both drought and irrigation caused individual plants to grow more between years. Population growth rates were typically higher with earlier flowering, but with phenological shifts of the magnitude that we observed, changes in λ were small. Furthermore, differences in λ between treatments were primarily attributable to increased growth, with only very small influences of phenology-driven changes to seed output on population growth. These results suggest that changing rainfall may influence local population dynamics of *L. utriculatum*, but not necessarily through changing flowering times and corresponding changes in seed production.

### Life history and demographic context

Our finding that population growth was only weakly affected by the shifts in flowering phenology we observed, particularly compared to changes in individual growth, align with life-history and demographic theories in two ways. First, long-lived plants commonly have population growth rates more sensitive to survival or growth than to reproduction (Franco & Silvertown 2004). Low sensitivity of λ to reproduction means that very large changes to reproductive output are needed to substantially alter the population growth rate. However, we observed phenological shifts that produced only small (typically <10%) changes in seed set. Instead, small changes to individual growth largely determined the ultimate response of λ to rainfall manipulation, due to the outsized sensitivity of λ to growth. In this sense, our results resemble those of Iler et al. (2019) who found that reduced survival had a larger influence on population growth of an alpine perennial than snowmelt-induced earlier flowing. In contrast, population dynamics of annual plants may shift more under climate change than for our perennial plant, not only because of the greater sensitivity of λ to reproduction in shorter-lived plants (Franco & Silvertown 2004), but also because annual plants often have greater phenological sensitivity to climate (König et al. 2018).

Second, negative vital rates correlations can stabilize population dynamics in the face of environmental variation (Villellas et al. 2015). In our study, we saw vital rates respond to rainfall manipulation in opposing directions. Namely, seed output declined in response to both rainfall manipulation treatments, but individual growth increased. These changes partially offset each other, damping the overall effect of changing rainfall on λ (Fig. 5). Opposing responses to changing rainfall also occurred for individual components of reproductive fitness (as seen by, e.g., Ehrlén & Münzbergova 2009): earlier flowering increased the probability of inflorescence failure but increased the number of seeds among the fraction of successful inflorescences, weakening the overall sensitivity of seed output to flowering phenology. It is not guaranteed, though, that plant populations will always show these negative correlations among vital rates. For example, Campbell (2019) found that all vital rates in an alpine perennial responded negatively to earlier snowmelt, producing a strongly negative response to the climate stressor.

### Phenological, physiological, and demographic responses to precipitation

Flowering time in *L. utriculatum* is sensitive to growing season moisture availability: decreased rainfall produced earlier flowering and increased rainfall delayed flowering in our multi-year manipulative experiment. Because the end of *L. utriculatum*’s growing season coincides with the onset of the summer dry season, earlier flowering with declining available moisture is consistent with drought escape and reproductive assurance (Kenney et al. 2014, Kooyers 2015). Additionally, earlier-flowering plants tended to produce slightly more seeds on average regardless of treatment, further suggesting it is important for *L. utriculatum* plants to synchronize reproduction with water availability. Temporal changes in pollinator behavior, composition, or effectiveness, an alternative mechanism for phenological relationships with reproductive success (Elzinga et al. 2007), seem unlikely given that pollen limitation appears to be uncommon at our site (Gielens et al. 2014). Given this sensitivity of flowering to moisture availability, it seems likely that flowering will initiate earlier in our study area as drier summers become more common, although this may be ameliorated somewhat by increasingly wet winters (CRVD 2017).

Yet, despite drought inducing earlier flowering and earlier flowering typically yielding more seeds, we did not find that drought increased seed output. This appears to be the result of direct, non-phenological effects of drought offsetting benefits of earlier flowering. This may be due to the common pattern of plants under moisture stress producing fewer flowers (Kuppler & Kotowska 2021) or lower quality pollen (Brunet et al. 2025). This result and the finding that changing individual growth dominated the changes in λ for both treatments demonstrate that phenological shifts are one of many important consequences influencing demography of plant populations under climate change. Recent studies have distinguished between phenological and non-phenological effects on individual components of fitness under climate change. Zettlemoyer et al. (2024) found that even *Silene acaulis* individuals that were able to shift their flowering times under warming and earlier snowmelt still produced fewer seeds, suggesting a climate change penalty independent of flowering time. Zhang et al. (2024) and Collins et al. (2025) compared phenological and non-phenological effects of warming on seed production in alpine and tundra, respectively, and found opposing results, with phenological and direct climate effects of similar magnitude in alpine plants but increased seed production in the tundra primarily attributable to phenological shifts, suggesting that the relative influences of these processes are likely context-dependent. We show that it is necessary to extend these results to consider effects on multiple vital rates, particularly those that most strongly influence λ or lifetime fitness. Thus, while shifting flowering times under climate change often increase fitness (Cleland et al. 2012), they may not be sufficient for persistence when weighed against concurrent negative fitness consequences of the climate stressor.

### The importance of individual growth

Increased individual growth under drought and decreased growth for small plants under irrigation defied our expectations that water deficit would impede growth. One possible explanation is that response of growth to precipitation was mediated by rainfall-induced changes to the competitive environment (Eskilinen & Harrison 2015). Prior work in our system found increasing grass and forb productivity with increased soil moisture (Smith & Williams 2023). Alleviating water stress often increases above-ground competition (Foxx & Fort 2019), potentially shading small plants and favoring more leaf area in plants with sufficient storage to produce it. Drought may correspondingly reduce aboveground competition, and *L. utriculatum* may be less sensitive to drought than other resident species due to its taproot or leaf morphology (Comas et al. 2013, Chitwood & Sinha 2016).

Changes in individual growth may also be due to changes in vegetative phenology (e.g., leaf-out timing), inviting future research. While there have been few studies examining the effects of changing flowering phenology under climate change and population dynamics in long-lived plants (Campbell 2019, Iler et al. 2019, Keller & Shea 2021), even fewer studies consider the role of vegetative phenology in perennials (Iler et al. 2021). While leaf-out timing tends to vary less than reproductive timing under climate change (Collins et al. 2021), given the high sensitivity of λ to survival and growth in long-lived plants, even modest shifts in vegetative phenology may affect population growth if they produce changes in resource acquisition, storage, and future growth. Earlier vegetative phenology *via* increased warming and/or earlier snowmelt has been shown to increase growth and productivity in some (Arft et al. 1999, Natali et al. 2012) but not all (Livensperger et al. 2016, Möhl et al. 2022) cases. Potential benefits of earlier leaf-out under decreased precipitation in our system may be at least partially negated by earlier senescence, considering drought typically favors early senescence in environments where the growing season terminates with a dry period (Kooyers 2015, Reed et al. 2019, Currier & Sala 2022).

### Conclusions

Phenology remains a key indicator of climate change and shifts in timing can be critically important for other biological processes, even as we found it was only marginally important for population growth in our study. Shifts in phenology can disrupt biotic interactions, such as pollination (Elzinga et al. 2007, Rafferty & Ives 2011) or frugivory (Deacy et al. 2017). Reproductive phenology also determines the temporal and environmental context for sexual reproduction, affecting evolutionary processes such as local adaptation (Hall & Willis 2006) or genetic differentiation *via* reproductive isolation (Hendry & Day 2005). Yet, population growth rates are a synthesis of several demographic processes, which may change or fluctuate with the same climate drivers that affect phenology (Williams et al. 2025). We offer two conclusions for consideration. First, we find little evidence suggesting that flowering phenology will influence population dynamics of a long-lived wildflower as it faces changes in precipitation. Second, we stress the importance of comparing the consequences of climate-driven phenological shifts for biological processes with the direct consequences of the climate stressor itself on said process. This distinction is crucial for accurately understanding mechanisms influencing species persistence and predicting the fates of plant populations.

## Acknowledgements

Field work took place on the traditional territory of Cowichan Tribes (Hul’qumi’num people). We thank the Nature Conservancy of Canada for site access and assistance, H. Bloom, S. Duncan, N. How, J. Johnson, D. Loughnan, E. McHugh, L. Smith, C. Trowbridge, and L. Worden for assistance with data collection, I. Banman for constructing the drought shelters, and L. White for constructing and maintaining the irrigation system. A. Angert, R. Bradley, J. Moutouama, and C. Urquhart provided helpful feedback on the manuscript. This research was supported by the Canada Foundation for Innovation, British Columbia Knowledge Development Fund, and a Natural Sciences and Engineering Research Council of Canada Discovery Grant to J.L.W. Analysis was assisted by Research Computing at Portland State University and its resources acquired through NSF grants 2019216 and 1624776.

