## Supporting Information for "Earlier flowering explains only a small part of experimental drought’s effects on a wildflower’s population growth"

### Supporting Methods and Materials

#### *Phenology and vital rate data and estimation*

Here we describe in more detail how we fit statistical models to estimate the effects of the rainfall manipulation experiment and phenology on vital rates of the perennial forb *Lomatium utriculatum*. In all candidate models, treatment (drought, irrigated, and unmanipulated control) was modeled as a fixed effect with the control treatment as the reference group. Inter-annual variation was modeled as a fixed effect of year for datasets with fewer than five years and as a random intercept where five or more years of data were available. We included year effects in all candidate models except for survival, where preliminary analysis suggested including random effects of year did not improve model fit. In all vital rate models, our candidate model set included models with two-way interactions (size-treatment and year-treatment effects), and we included size-year effects in models fit with yearly fixed effects. To reduce the number of candidate models tested, we did not test for three-way (size-year-treatment) interaction effects. See Table S1 for a full list of vital rates and dataset extents used for each.

We modeled survival with logistic GLMMs. Because we imputed survival for individuals not detected in a census year but found alive the following year, we did not include the final census year in our survival models, as we could not determine whether plants not observed were dead or undetected for other reasons (e.g., dormancy). We modeled growth, conditioned on survival, as a mixed effects model with a Gaussian-distributed error term. To quantify trade-offs between phenology in year $t$ and performance in year $t+1$, we also tested for effects of flowering time in the prior year on both survival and growth by fitting models with the subset of observations 2021-2024 with recorded flowering dates.

To estimate inflorescence production, we fit models for the number of inflorescences as zero-inflated GLMMs with a truncated Poisson distribution for the conditional response. All plants in the census for which size was recorded were included, with one observation per plant per year. The zero-inflation portion of the model gives the probability of producing zero inflorescences; correspondingly, one minus the prediction of the zero-inflation component is the probability of flowering. The conditional portion of the model gives the number of inflorescences per flowering plant. The product of the predicted probability of flowering and the number of inflorescences per flowering plant is the predicted number of inflorescences per plant. Fifty percent of observed flowering plants produced one inflorescence and the maximum number of inflorescences observed on a plant was 11.

To estimate seed production per inflorescence, we fit zero-inflated GLMMs with a negative binomial distribution for the conditional response. All inflorescences in the dataset for which seed set and plant-level phenology could be estimated were included, with inflorescence as the unit of observation. The zero-inflation portion of the model estimates the probability of inflorescences mortality before setting seed, e.g., due to grazing or abortion, such that one minus the prediction of the zero-inflation component gives the probability of an inflorescence surviving to produce seed (inflorescence “success”). The conditional portion of the model gives the number of seeds produced per successful inflorescence. The product of the inflorescence success probability and number of seeds per successful inflorescence gives the predicted number of seeds per inflorescence. For the inflorescence success model component, we also tested for an effect of the number of inflorescences produced by the plant.

We designated observed plants as new recruits (i.e., established seedlings in their first growing season) if they met each of the following conditions: (1) observation was the first sighting of the plant, (2) observation occurred in 2018 or later (i.e., after the first two years of demographic surveys in the plots), (3) plant had only one leaf, and (4) plant was not flowering in the year of observation. This filtering process left 318 observations across seven years. We fit models with size of the recruit as the response with a Gaussian error term. We tested for effects of rainfall treatment, including annually-varying treatment effects.

We ran a one-year *in situ* seed addition study to inform our modeling of seedling establishment rates. However, because of the lack of temporal replication we did not use these estimates in our kernels; instead, we numerically solved for a seedling establishment rate that fixed λ=1 for the control treatment and used this value (~0.0046) for all kernels. We summarize the experiment and its results here for comparison with the establishment rate that we used in analysis. In the seed addition study, seeds were collected in June 2021 shortly before addition; this timing corresponds to the natural seed dispersal timing at the field site. Eight 10x10 $cm^2$ subplots were established in each plot and 30 *L. utriculatum* seeds were added to two subplots per plot. For a related project assessing seed predation, four subplots were covered by wire mesh predator exclusion cages. One plot only had four subplots (uncaged) established due to extremely dense vegetation, leaving 116 observations. We returned to count the number of germinants present in 2022 and 2023. No additional germinants were observed in 2023, suggesting no seed dormancy at this site. We estimated the mean number of germinants observed per subplot with a GLMM with a negative binomial response, including a fixed effect for *L. utriculatum* addition and a random intercept for each plot. There was no difference in germinant presence between caged and uncaged subunits, nor were there differences in germinant presence among treatments. Our final model predicted 0.0135 (95% confidence interval: [0.002, 0.081]) seedlings per plot without addition (from natural seed rain) and 0.2850 (CI: [0.069, 1.141]) seedlings per addition plot. We then constructed the following system of equations:

$$y_0 = px$$
$$y_1 = p(x + 30),$$

where $y_0$ and $y_1$ are the respective number of seedlings in plots without or with seeds added, $p$ is the per-seed probability of establishment, and $x$ is the background number of seeds present per plot. Substituting in our observed values and solving for $p$ gives an estimated germination probability in 2022 of 0.0090 and 95% confidence interval (0.002, 0.035). The germination rate used in our kernels (~0.0046) lies within this interval.

*Integral projection model implementation*

We discretized plant size into bins of width 0.1, with respective lower and upper bounds 0.045 and 6.05 such that when using the midpoint rule vital rate models were evaluated over the range 0.5 to 6.0; these bounds were chosen because they covered the entire range of plant sizes observed in our surveys. As such, plant size was represented as a vector with 57 elements and kernels were estimated as 57 x 57 matrices. These bounds produced few "evictions" of small and large individuals (potential error in λ < 10^(-5); Williams et al. 2012).

We estimated kernels by evaluating model predictions across our size range, using mean flowering date and treatment effects to generate phenology- and treatment-specific kernels. Model predictions were generated with all random intercepts set to zero, i.e., for the "average" plot, plant, and year where year was modeled as a random effect. In models where annual variation was estimated with fixed effects, estimates were generated on the linear scale (i.e., estimated before being back-transformed through the link function) for each year of observation used to fit the model, averaged across all years, and then transformed through the link function to produce a mean estimate. The one exception is that estimates of inflorescence success in 2021 were removed from the inflorescence success average, as the limited sample size in that year meant that the estimate of the inflorescence failure rate was overly influenced by outliers and produced unrealistic inflorescence failure rates for very large plants.

*Life Table Response Experiment*

Our full derivation is provided at the end of the Supporting Information (see Additional Supporting Information). Sensitivities to each vital rate parameter were estimated by re-estimating kernels with perturbed vital rates and estimating the change in λ (approach 5 in g17). For each LTRE between two matrices, $K_0$ and $K_1$, we estimated sensitivity to each parameter at both $K_0$ and $K_1$ and averaged the two sensitivities to approximate the midpoint sensitivity.

In our first-order LTRE (i.e., one without second-order terms for phenology-treatment interactions), the distinct LTRE contributions of treatment and phenology can be estimated in two separate ways. One set of estimates has the treatment effects (ψ terms) estimated on the flowering date observed in controls and the phenology effects (φ terms) estimated within the non-control treatments; the other estimates have phenology effects estimated within the control treatment and the direct treatment effects estimated on the flowering dates observed under drought/irrigation. Visually, the first of these is represented in Figure 1 of the main text. We refer to this second, alternative set of results as the "mirrored" analysis. We present the first set of results as Fig. 5 in the main text (with ψ terms estimated on the control mean flowering date and φ terms estimated within the drought/irrigation treatments, *sensu* Fig. 1). Because Δλ is close to linear with respect to flowering dates, the mirrored results were nearly identical to the main results (Table S19).

### Supporting Results

*Phenology analysis results*

The best-supported phenology model featured a fixed effect of treatment, with advancing flowering under drought and delays in flowering under irrigation (Table S3). A model with annually varying treatment effects that similarly showed consistently advanced flowering under drought and delayed flowering under irrigation, but only in some years (Fig. S1), had slightly lower AIC (model with interaction terms: ΔAIC= 0.98; Table S2), but since it had more parameters, this model did not have enough support to adopt as a final model by our selection criteria. Models using each plant's first flowering day (the earliest flowering date among all inflorescences produced by a plant in a year, rather than the average across inflorescences) as a response produced qualitatively similar results: a multi-day advance in flowering under drought, a non-significant delay under irrigation, and no support for an interaction (Tables S4-5).

Rainfall manipulation influencing phenology had the potential to limit our ability to identify separate contributions of treatment and phenology to variation in vital rates. We evaluated this possibility by examining the variance in phenology explained by the rainfall manipulation treatment; if rainfall treatment explained a substantial portion of overall variance in flowering times, identifying separate contributions of rainfall and flowering would be impossible. We found that rainfall treatment increased the pseudo-$R^2$ from 0.27 in a mixed effects model with only fixed effects of year to 0.32 (equations 26-27, Nakagawa & Schielzeth 2013), i.e., adding treatment to the model only explained 7% of unexplained variance after accounting for annual variation. This can be illustrated by comparing the variance terms associated with the model random intercepts. A model without treatment effects had among-plot standard deviation of 2.5 days; including treatment effects reduced this to 1.4 days (a 67% reduction in variance). However, the model's other variance components were larger than the among-plot variance (within-plot standard deviation: 3.9 days, within-plant standard deviation 6.7 days). Thus, while treatment effects did explain some variation in flowering times, there was considerable residual variance in plant mean flowering dates that treatment could not explain. This non-treatment variance in flowering times allowed us to separately identify treatment effects on vital rates and phenological effects on vital rates that were not attributable to treatment.

*Vital rate model results*

We found no evidence that rainfall manipulation treatment affected survival (Table S6). Our final model featured only a fixed effect of size, with higher survival for larger plants (Table S7). Likewise, there was no evidence of a trade-off between survival and phenology among plants flowering in the prior year (Table S8). Rainfall manipulation had size-dependent effects on growth (Tables S9-10). In particular, on average larger plants under irrigation and smaller- and medium-sized plants under drought grew more than their counterparts in control conditions (Fig. S2). We found evidence of a trade-off with phenology such that after statistically controlling for treatment, plants that flowered early in the prior year grew slightly more than their later-flowering counterparts (Tables S11-12).

Plants under irrigation were less likely to flower on average compared to plants under control conditions (Tables S13-14, Fig. S3a). Among those plants that did flower, rainfall manipulation did not affect the number of inflorescences produced (Table S14, Fig. S3b) or the probability of inflorescences surviving to produce seed (Tables S15-16, Fig. S3d). Inflorescence success tended to decrease as the number of inflorescences a plant produced increased (Table S16), which would be consistent with plants producing inflorescences later in the season if earlier-season inflorescences fail due to, e.g., grazing or abortion. Treatment had size-dependent effects on the number of seeds produced per successful inflorescence: plants under drought typically produced fewer seeds regardless of size, but smaller plants under irrigation tended to produce more seeds than plants in control conditions (Fig. S3e). The combined effects of these direct treatment effects was to slightly reduce seed production in both manipulation treatments, particularly among larger plants (Fig. S3g,h). We also found that mean timing of flowering was associated with both increased inflorescence success and number of seeds produced per successful inflorescence, but these effects acted in opposing directions (Table S16). We did not find evidence of quadratic relationships between flowering time and either component of fitness (Table S15).

Rainfall manipulation also affected the observed size of new recruits (Table S17). New recruits observed under drought tended to be ~20% larger than plants in control conditions (Table S18).

*Life Table Response Experiment*

Our analysis produced similar results when using different establishment rates. For both treatments, $\lambda$ was larger in treatments compared to controls and tended to increase with higher establishment rates, although uncertainty in $\Delta\lambda$ also increased with increasing establishment probability (Fig. S4). Likewise, the magnitudes of the individual LTRE contributions all increased in magnitude with the establishment rate, but the pattern that rainfall treatment effects dominate with positive effects acting through growth and negative effects acting through reproduction remained consistent (Fig. S5). Main and mirrored LTRE contributions for individual vital rates were highly correlated with each other and produced qualitatively identical results (Table S19).

**Additional Supporting Information**

*LTRE derivation*

Let *Y* be a random variable defining a vital rate that is a function of size, $z$, estimated with generalized linear models. All vital rate model coefficients can be classified as either shifting the intercept or shifting the slope (i.e., size-independent or size-dependent). Summing together all respective intercept and slope terms evaluated at controls and writing them as $a_0^{(cont)}$ and $a_1^{(cont)}$, the expectation of the vital rate in controls can be written as

$$E[Y^{(cont)}|z] \;=\; g(a_0^{(cont)} + a_1^{(cont)}z),$$

for link function *g*. The right-hand side of the above expression is equivalent to the prediction from the vital rate's GLMM evaluated for controls.

Our vital rate models estimated direct rainfall treatment effects using categorical variables, with estimates quantifying the marginal effect of drought or irrigation treatment on the vital rate's intercept or slope relative to the controls. Effects of phenological shifts were estimated as continuous (i.e., per-day) effects shifting the intercept. We did not test for size-dependent (i.e., slope-shifting) phenological effects in our analysis. The vital rate predictions for drought and irrigation, including both direct treatment and phenological effects, are thus

$$E[Y^{(dro)}|z] \;=\; g([a_0^{(cont)}+\beta_0^{(dro)} + \gamma_0\ \bar{x}_{dro}] \;+ [a_1^{(cont)}+\beta_1^{(dro)}]z),$$

and

$$E[Y^{(irr)}|z] \;=\; g([a_0^{(cont)}+\beta_0^{(irr)} + \gamma_0\ \bar{x}_{irr}] \;+ [a_1^{(cont)}+\beta_1^{(irr)}]z),$$

where $\beta_0^{(dro)}$ ($\beta_0^{(irr)}$) is the marginal effect of drought (irrigation) on the vital rate intercept, $\beta_1^{(dro)}$ ($\beta_1^{(irr)}$) is the marginal effect of drought (irrigation) on the vital rate slope, $\gamma_0$ is the per-day effect of the phenological shift on the vital rate intercept, and $\bar{x}_{dro}$ ($\bar{x}_{irr}$) is the mean phenological shift associated with drought (irrigation), measured in days. The β and γ terms are coefficients from the vital rate models and the *x* terms were estimated from the model of treatment effects on phenology. Vital rates with no differences among treatments or covariance with phenology have β and γ equal to zero.

The effects of drought on λ acting through vital rate *Y* can then be expressed as:

$$\beta_0^{(dro)}\frac{\partial\lambda}{\partial a_0} + \gamma_0\underline{x}_{dro}\frac{\partial\lambda}{\partial a_0} + \beta_1^{(dro)}\frac{\partial\lambda}{\partial a_1},$$

where the three terms in the sum are the respective contributions of a direct treatment-driven shift to the vital rate model intercept, the phenology-driven shift to the vital rate model intercept, and the direct treatment-driven shift to the vital rate model slope. A similar expression gives the effects of irrigation on λ acting through vital rate *Y*. Drought and irrigation effects acting through other vital rates are modeled accordingly. For vital rate *Y*, we can express the respective direct treatment and phenological contributions to Δλ as

$$\psi_Y^{(dro)} = \beta_0^{(dro)}\frac{\partial\lambda}{\partial a_0} + \beta_1^{(dro)}\frac{\partial\lambda}{\partial a_1}, \text{ and}$$

$$\varphi_Y^{(dro)} = \gamma_0\bar{x}_{dro}\frac{\partial\lambda}{\partial a_0},$$

with similar expressions giving the respective contributions to Δλ under irrigation.

If the set of vital rates is denoted as *V,* then the respective cumulative contributions of direct treatment effects and phenological shifts on Δλ are

$$\psi^{(dro)} = \sum_{v \in V} \psi_v^{(dro)} \text{ and}$$

$$\varphi^{(dro)} = \sum_{v \in V} \varphi_v^{(dro)},$$

such that

$$\Delta\lambda^{(dro)} \approx \psi^{(dro)} + \varphi^{(dro)}.$$

The effects of irrigation on λ are defined similarly.

**Supporting Figures**

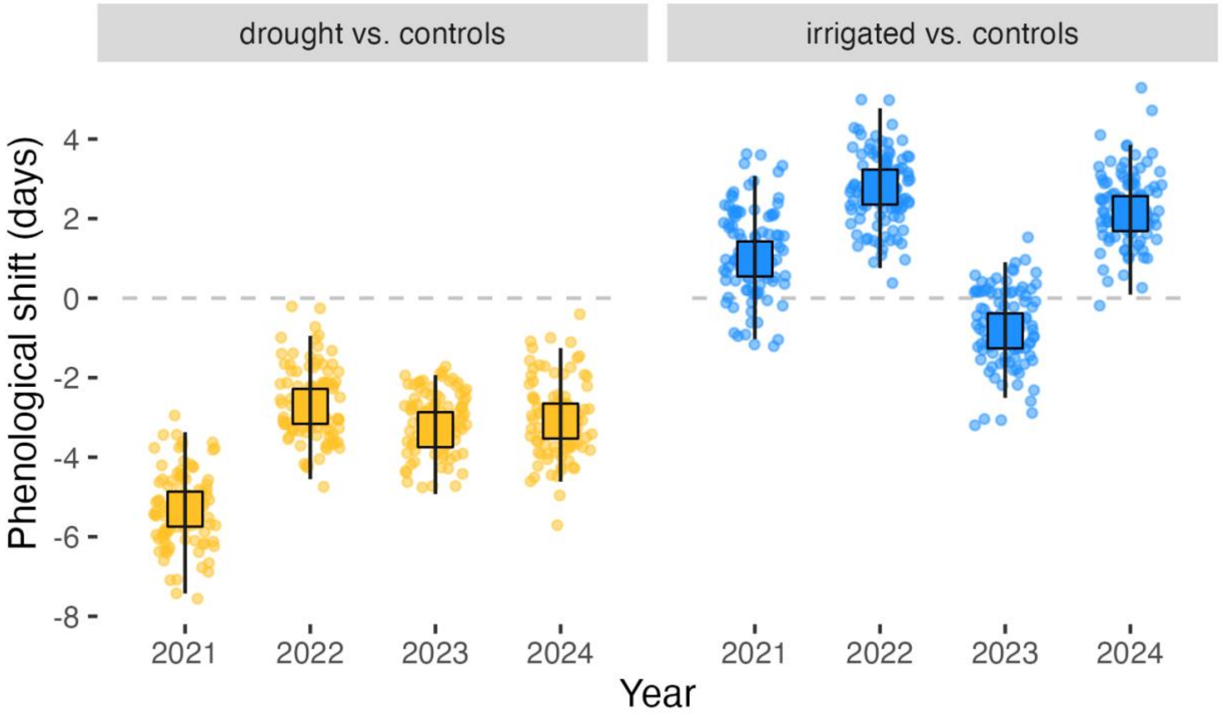


**Fig. S1** Estimated differences in phenology between rainfall manipulation treatments and controls, predicted by a less-supported model with annually-varying treatment effects. Squares give mean annual differences between treatment and controls, segments are 95% bootstrapped confidence intervals of the difference, and small circles are a sample of bootstrapped treatment effects from refit models. Grey dashed line at zero days denotes zero difference between treatment and control.

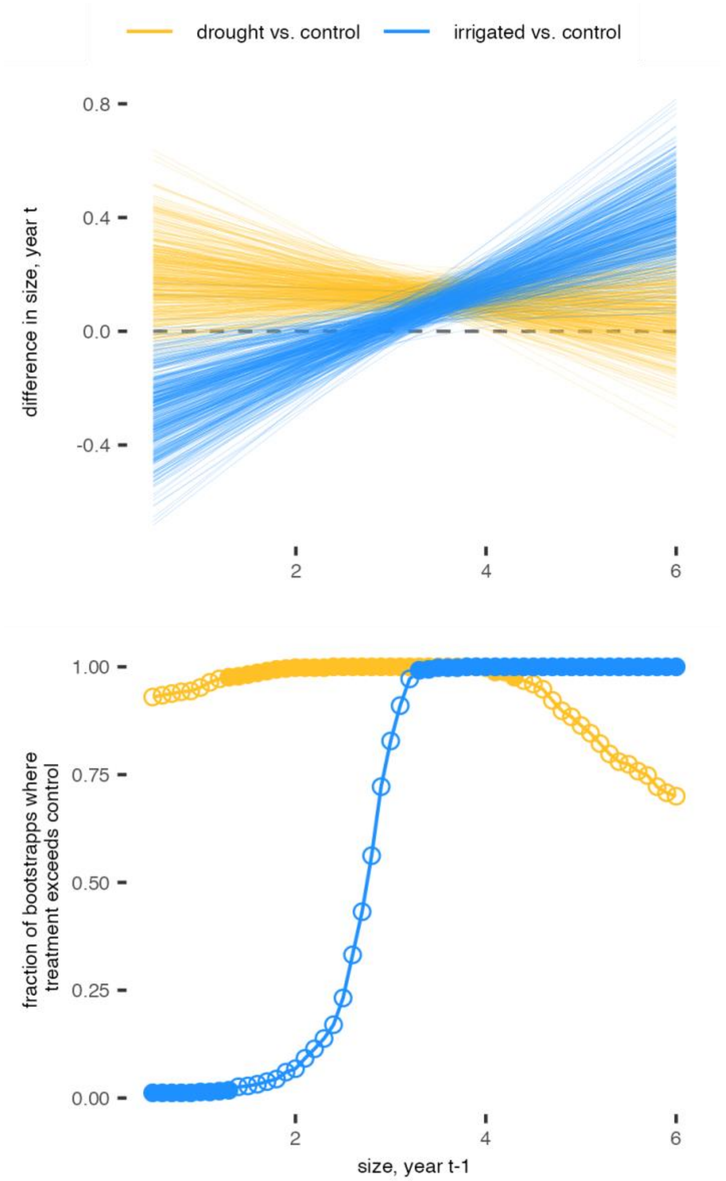


**Fig. S2** Estimated differences in mean annual individual growth between treatments and controls, re-estimated with 500 bootstrapped datasets. Top panel gives the difference between treatment and control for either treatment for each bootstrapped model. Small plants in irrigated plots grew less than those in controls, but small plants in drought plots grew more; the converse was true for large plants. Bottom panel gives the fraction of bootstraps where the given treatment has higher greater growth than the controls at the given size. Filled circles represent differences in growth that are significant at the $\alpha$=0.05 level (i.e., fewer than 2.5% or greater than 97.5% of bootstrapped estimates exceeding controls at the given size value).

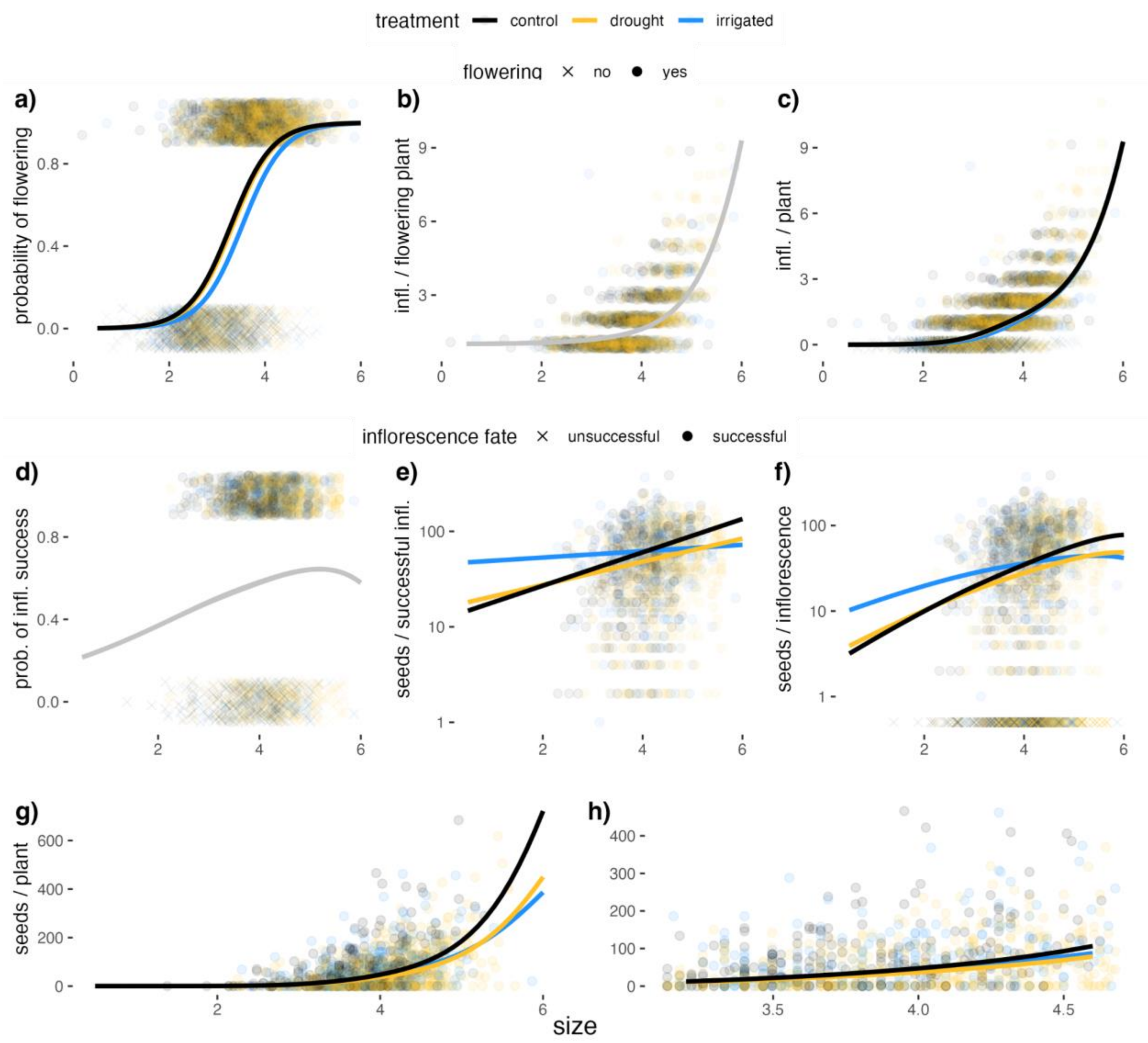


**Fig. S3** Summary of reproductive vital rate relationships with plant size and rainfall treatment. a) Probability of flowering; b) inflorescences per flowering plant; c) inflorescences per plant (i.e., product of components in panels a and b); d) probability of inflorescence surviving to produce non-zero number of seeds (i.e., inflorescence success); e) seeds per successful inflorescence; f) seeds per inflorescence (i.e., product of components in panels d and e); g) total number of seeds per plant; h) total number of seeds per plant with size range subsetted to between the 10th and 90th percentiles for observed size of flowering plants. Gray curves in panels b) and d) indicate no differences among treatments. Points in panel a-c and g-h each represent one plant; points in panel d-f each represent one inflorescence. Estimates in panels d-f are for plants with mean flowering date equal to the mean flowering date observed in controls.

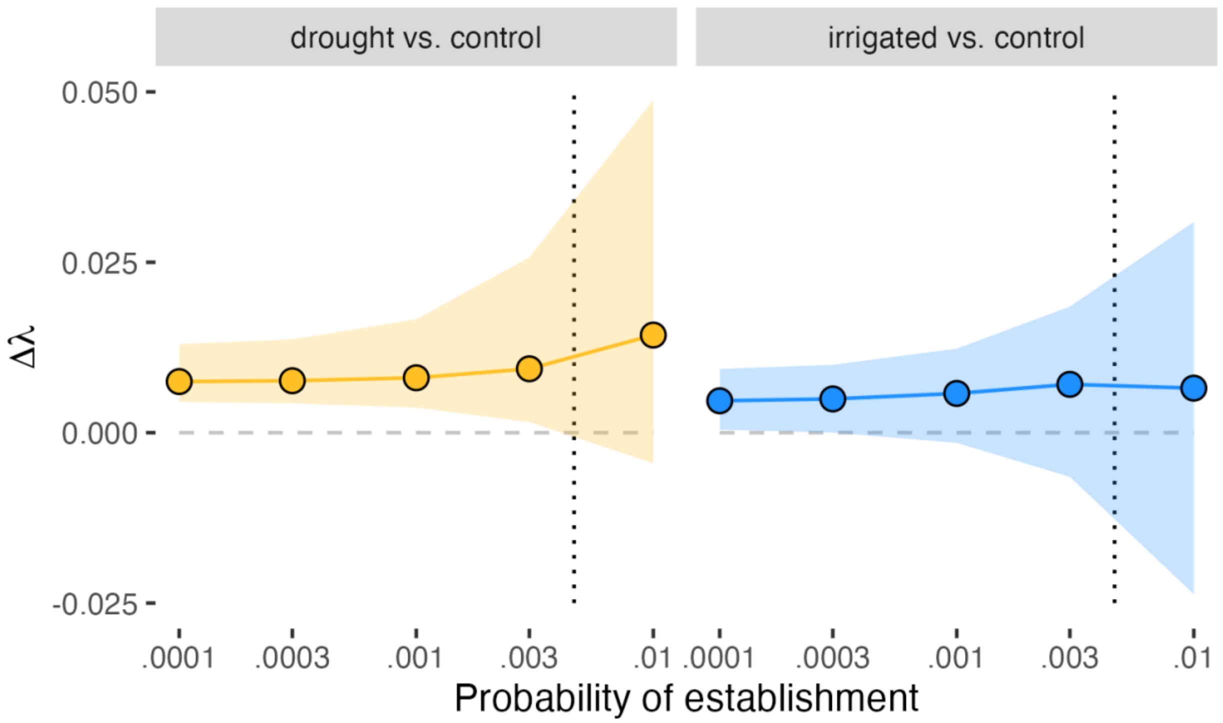


**Fig. S4.** Estimates of Δλ for treatments relative to controls, estimated across establishment probabilities. Shaded regions are 95% bootstrapped confidence intervals. Dotted vertical line indicates the establishment rate used in analysis (~0.0046).

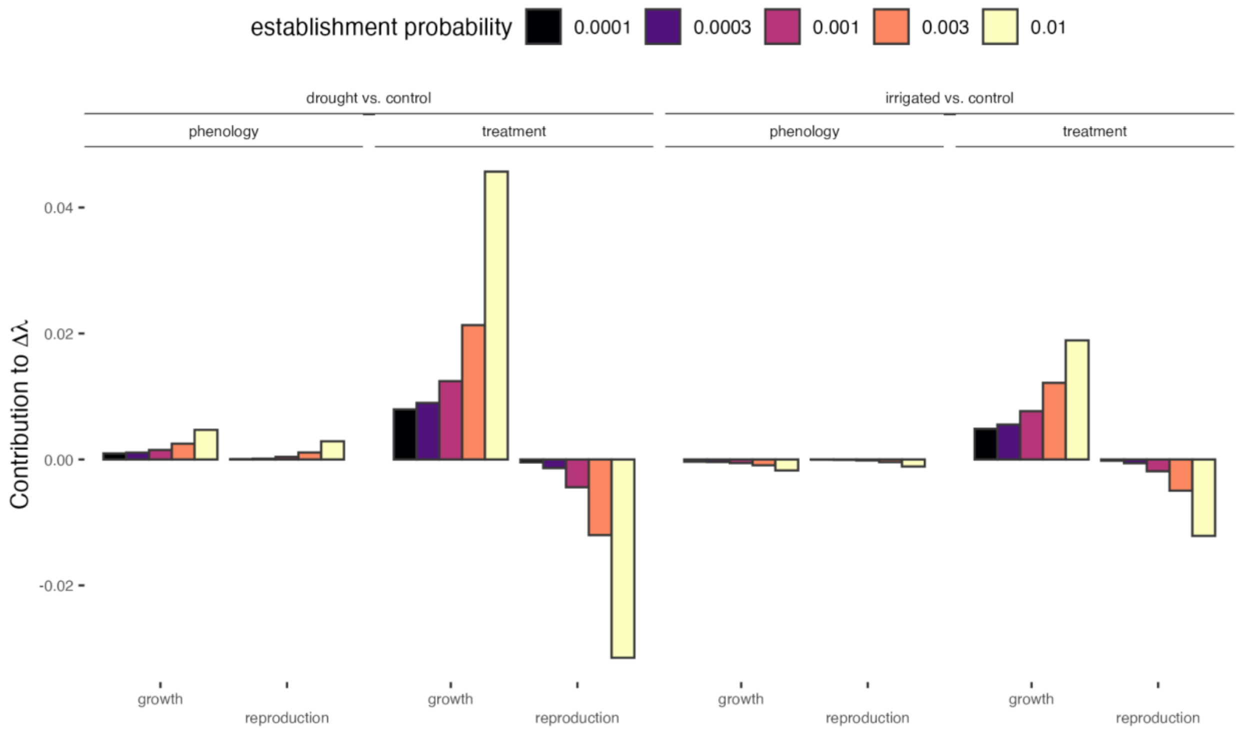


**Fig. S5.** LTRE contributions (grouped by growth/reproduction and phenology/treatment-driven) estimated with different establishment probabilities.

**Supporting Tables**

| Vital rate | Years/transitions observed | Number of observations | Number of unique plants |
|---|---|---|---|
| Survival[a] | 6 | 3087 | 936 |
| Growth[b] | 7 | 2970 | 843 |
| Probability of flowering | 8 | 4734 | 1056 |
| Number of inflorescences | 8 | 4734 | 1056 |
| Probability of inflorescence success | 4 | 2372 | 564 |
| Seeds per successful inflorescence | 4 | 2372 | 564 |
| Establishment probability[c] | N/A | N/A | N/A |
| Recruit size | 7 | 318 | 318 |

**Table S1:** Description of vital rates, years of data included in model estimates, number of observations in dataset, and number of unique plants included in dataset.

[a] - Survival-phenology trade-off evaluated with two annual transitions of data (2021-2023), 632 observations of 450 plants
[b] - Growth-phenology trade-off evaluated with three annual transitions of data (2021-2024), 840 observations of 476 plants
[c] - Probability of seed transitioning into recruit; see Supporting Methods for estimation details

| Model structure | *p* | AIC | ΔAIC |
|---|---|---|---|
| `phen ~ Year + (1 | Plot / plantid)` | 7 | 9263.84 | 8.22 |
| **`phen ~ trt + Year + (1 | Plot / plantid)`** | **9** | **9256.60** | **0.98** |
| `phen ~ trt + (1 | Year) + (1 | Plot / plantid)` | 7 | 9274.53 | 18.91 |
| `phen ~ trt * Year + (1 | Plot / plantid)` | 15 | 9255.62 | 0.00 |

**Table S2:** Models fit to select the best phenology model. Coefficients: `trt` is a categorical treatment, and `Year` (categorical variable) is year of census. Asterisk (`*`) denotes both additive terms and an interaction term and parenthesized terms are model random effects. *p* denotes the number of parameters estimated. Bolded model is the final model used to estimate phenology effects.

| Effect type | Coefficient name | Estimate | Std. error |
|---|---|---|---|
| Fixed | `Intercept` | 118.956 | 1.016 |
| | `trtdrought` | -3.360 | 1.236 |
| | `trtirrigated` | 1.300 | 1.306 |
| | `Year2021` | 5.098 | 0.600 |
| | `Year2022` | 12.280 | 0.493 |
| | `Year2023` | 9.552 | 0.533 |
| Random | `Plot` | 1.424 | - |
| | `plantid:Plot` | 3.891 | - |
| | `Residual` | 6.696 | - |

**Table S3:** Model coefficients in final phenology model used in analysis. Reference group (`Intercept`) is the control treatment in year 2024. Estimates for random effects are the standard deviations associated with the random intercepts.

| Model structure | *p* | AIC | ΔAIC |
|---|---|---|---|
| `phen.first ~ Year + (1 | Plot / plantid)` | 7 | 10009.21 | 9.40 |
| **`phen.first ~ trt + Year + (1 | Plot / plantid)`** | **9** | **9999.81** | **0.00** |
| `phen.first ~ trt + (1 | Year) + (1 | Plot / plantid)` | 7 | 10016.27 | 16.45 |
| `phen.first ~ trt * Year + (1 | Plot / plantid)` | 15 | 10008.67 | 8.86 |

**Table S4:** Models fit to evaluate treatment effects on a plant's first day of flowering (`phen.first`). Coefficients: `trt` is a categorical treatment, and `Year` (categorical variable) is year of census. Asterisk (`*`) denotes both additive terms and an interaction term and parenthesized terms are model random effects. *p* denotes the number of parameters estimated. Bolded model is the best supported model of those tested.

| Effect type | Coefficient name | Estimate | Std. error |
|---|---|---|---|
| Fixed | `Intercept` | 114.355 | 0.945 |
| | `trtdrought` | -3.969 | 1.078 |
| | `trtirrigated` | 0.801 | 1.174 |
| | `Year2021` | 9.682 | 0.797 |
| | `Year2022` | 11.301 | 0.657 |
| | `Year2023` | 10.971 | 0.709 |
| Random | `Plot` | 0.929 | - |
| | `plantid:Plot` | 4.935 | - |
| | `Residual` | 8.947 | - |

**Table S5:** Model coefficients in best supported model predicting first flowering date. Reference group (`Intercept`) is the control treatment in year 2024.

| Model structure | *p* | AIC | ΔAIC |
|---|---|---|---|
| ~ (1 \| Plot) | 2 | 1829.01 | 51.82 |
| **~ size.prev + (1 \| Plot)** | **3** | **1777.19** | **0.00** |
| ~ (1 \| Year) + (1 \| Plot) | 3 | 1827.44 | 50.25 |
| ~ size.prev + (1 \| Year) + (1 \| Plot) | 4 | 1779.19 | 2.00 |
| ~ trt + size.prev + (1 \| Plot) | 5 | 1780.23 | 3.05 |
| ~ trt * size.prev + (1 \| Plot) | 7 | 1782.45 | 5.26 |

**Table S6:** Models fit to select the best model of survival. Coefficients: `size.prev` is size of plant in the year prior to the census (i.e., size before survival/mortality), `trt` is a categorical treatment, and `Year` is year of census. Asterisk (`*`) denotes both additive terms and an interaction term and parenthesized terms are model random effects. *p* denotes the number of parameters estimated. Bolded model is the final model used to estimate kernels.

| Effect type | Coefficient name | Estimate | Std. error |
|---|---|---|---|
| Fixed | `Intercept` | 0.474 | 0.100 |
| | `size.prev` | 0.632 | 0.000 |
| Random | `Plot` | 0.332 | - |

**Table S7:** Model coefficients in final survival model used in kernels.

| Model structure | *p* | AIC | ΔAIC |
|---|---|---|---|
| **~ size.prev + (1 \| Plot)** | **3** | **221.06** | **0.00** |
| ~ Year + size.prev + (1 \| Plot) | 4 | 222.05 | 0.99 |
| ~ trt + size.prev + (1 \| Plot) | 5 | 224.71 | 3.65 |
| ~ phen + size.prev (1 \| Plot) | 4 | 222.37 | 1.31 |
| ~ phen^2 + size.prev + (1 \| Plot) | 5 | 223.64 | 2.58 |

**Table S8:** Models fit to test for effects of prior year's flowering phenology on survival. Coefficients: `size.prev` is size of plant in the year prior to the census (i.e., size before survival/mortality), `trt` is a categorical treatment, `Year` is year of census, `phen` is the mean-centered flowering date of the plant, and parenthesized terms are model random effects. Superscript/caret (`^2`) denotes both a linear and quadratic term for the variable in question.

| Model structure | *p* | AIC | ΔAIC |
|---|---|---|---|
| `~ size.prev + (1 | Plot / plantid)` | 5 | 6281.60 | 610.88 |
| `~ size.prev + (1 | Year) + (1 | Plot / plantid)` | 6 | 5786.13 | 115.42 |
| `~ size.prev + trt + (1 | Year) + (1 | Plot / plantid)` | 8 | 5788.98 | 118.26 |
| `~ size.prev + trt + (1 | trt:Year) + (1 | Year) + (1 | Plot / plantid)` | 9 | 5681.94 | 11.22 |
| **`~ size.prev * trt + (1 | trt:Year) + (1 | Year) + (1 | Plot / plantid)`** | **11** | **5670.72** | **0.00** |
| `~ size.prev * trt + (1 | Year) + (1 | Plot / plantid)` | 10 | 5777.63 | 106.91 |
| `~ size.prev + (1 | trt:Year) + (1 | Year) + (1 | Plot / plantid)` | 8 | 5778.57 | 107.86 |

**Table S9:** Models fit to select the best model of growth (main model). Coefficients: `size.prev` is size of plant in the year prior to the census, `trt` is a categorical treatment, `Year` is year of census/measurement. Asterisk (`*`) denotes both additive terms and an interaction term, colon (`:`) denotes only the interaction term, and parenthesized terms are model random effects. *p* denotes the number of parameters estimated. Bolded model is the final model used to estimate growth in kernels.

| Effect type | Coefficient name | Estimate | Std. error |
|---|---|---|---|
| Fixed | `Intercept` | 2.335 | 0.209 |
| | `size.prev` | 0.286 | 0.039 |
| | `trtdrought` | 0.207 | 0.225 |
| | `trtirrigated` | -0.348 | 0.228 |
| | `size.prev:trtdrought` | -0.024 | 0.043 |
| | `size.prev:trtirrigated` | 0.126 | 0.045 |
| Random | `Year` | 0.272 | - |
| | `trt:Year` | 0.178 | - |
| | `Plot` | 0.195 | - |
| | `plantid:Plot` | 0.202 | - |
| | `Residual` | 0.588 | - |

**Table S10:** Model coefficients for final model of growth used in kernels. Here, `size.prev` corresponds to a continuous variable (see previous table for definition) and `trt` corresponds to categorical variables with control as the reference group (i.e., term gives shift in intercept or slope for a treatment relative to the control). The `Residual` random effect is the Gaussian error term, i.e., within-individual level standard deviation after accounting for all other model sources of variation.

| Model structure | *p* | AIC | ΔAIC |
|---|---|---|---|
| `~ size.prev + Year + (1 | Plot / plantid)` | 7 | 1490.68 | 37.17 |
| `~ size.prev * Year + (1 | Plot / plantid)` | 9 | 1483.57 | 30.06 |
| `~ trt + size.prev * Year + (1 | Plot / plantid)` | 11 | 1485.13 | 31.62 |
| `~ trt*Year + size.prev*Year + (1 | Plot / plantid)` | 15 | 1463.84 | 10.33 |
| `~ size.prev * trt + size.prev * Year + (1 | Plot / plantid)` | 13 | 1488.27 | 34.76 |
| **`~ phen + trt * Year + size.prev * Year + (1 | Plot / plantid)`** | **16** | **1453.51** | **0.00** |
| `~ phen^2 + trt * Year + size.prev * Year + (1 | Plot / plantid)` | 17 | 1455.20 | 1.69 |
| `~ phen*Year + trt * Year + size.prev * Year + (1 | Plot / plantid)` | 18 | 1456.87 | 3.36 |

**Table S11:** Models fit to test for effects of prior year's flowering phenology on growth. Coefficients: `size.prev` is size of plant in the year prior to the census (i.e., size before survival/mortality), `trt` is a categorical treatment, `Year` is year of census, and `phen` is the mean-centered flower emergence date for the plant. Asterisk (`*`) denotes both additive terms and an interaction term, colon (`:`) denotes only the interaction term, and parenthesized terms are model random effects. *p* denotes the number of parameters estimated. Bolded model is the best-supported model; the `phen` estimate from this model is used in kernels.

| Effect type | Coefficient name | Estimate | Std. error |
|---|---|---:|---:|
| Fixed | Intercept | 2.686 | 0.284 |
| | size.prev | 0.293 | 0.069 |
| | Year2022 | -0.354 | 0.322 |
| | Year2023 | -1.415 | 0.361 |
| | trtdrought | 0.526 | 0.170 |
| | trtirrigated | 0.364 | 0.175 |
| | phen | -0.009 | 0.002 |
| | size.prev:Year2022 | 0.137 | 0.084 |
| | size.prev:Year2023 | 0.277 | 0.091 |
| | trtdrought:Year2022 | -0.562 | 0.120 |
| | trtdrought:Year2023 | -0.187 | 0.133 |
| | trtirrigated:Year2022 | -0.283 | 0.132 |
| | trtirrigated:Year2023 | -0.270 | 0.143 |
| Random | Plot | 0.185 | - |
| | plantid:Plot | 0.000 | - |
| | Residual | 0.556 | - |

**Table S12:** Model coefficients for best-supported model in testing for phenology-growth relationships. Here, `trt` corresponds to categorical variables with control as the reference group (i.e., term gives shift in intercept or slope for a treatment relative to the control), and `phen` is a continuous variable.

| Model structures | *p* | AIC | ΔAIC |
|---|---|---|---|
| ZI: ~ (1 \| Year) + (1 \| Plot / plantid)<br>Cond: ~ (1 \| Year) + (1 \| Plot / plantid) | 8 | 10255.83 | 1738.69 |
| ZI: ~ size + (1 \| Year) + (1 \| Plot / plantid)<br>Cond: ~ (1 \| Year) + (1 \| Plot / plantid) | 9 | 9235.36 | 718.22 |
| ZI: ~ (1 \| Year) + (1 \| Plot / plantid)<br>Cond: ~ size + (1 \| Year) + (1 \| Plot / plantid) | 9 | 9544.59 | 1027.22 |
| ZI: ~ size + (1 \| Year) + (1 \| Plot / plantid)<br>Cond: ~ size + (1 \| Year) + (1 \| Plot / plantid) | 10 | 8524.12 | 6.98 |
| ZI: ~ trt + size + (1 \| Year) + (1 \| Plot / plantid)<br>Cond: ~ size + (1 \| Year) + (1 \| Plot / plantid) | 12 | 8527.00 | 9.86 |
| ZI: ~ size + (1 \| Year) + (1 \| Plot / plantid)<br>Cond: ~ trt + size + (1 \| Year) + (1 \| Plot / plantid) | 12 | 8527.88 | 10.74 |
| ZI: ~ trt * size + (1 \| Year) + (1 \| Plot / plantid)<br>Cond: ~ size + (1 \| Year) + (1 \| Plot / plantid) | 14 | 8527.73 | 10.59 |
| ZI: ~ size + (1 \| Year) + (1 \| Plot / plantid)<br>Cond: ~ trt * size + (1 \| Year) + (1 \| Plot / plantid) | 14 | 8531.28 | 14.14 |
| **ZI: ~ trt + size + (1 \| Year) + (1 \| Year:trt) + (1 \| Plot / plantid)<br>Cond: ~ size + (1 \| Year) + (1 \| Plot / plantid)** | **13** | **8517.14** | **0.00** |
| ZI: ~ size + (1 \| Year) + (1 \| Plot / plantid)<br>Cond: ~ trt + size + (1 \| Year) + (1 \| Year:trt) + (1 \| Plot / plantid) | 13 | 8526.27 | 9.13 |
| ZI: ~ trt + size + (1 \| Year) + (1 \| Year:trt) + (1 \| Plot / plantid)<br>Cond: ~ trt + size + (1 \| Year) + (1 \| Year:trt) + (1 \| Plot / plantid) | 16 | 8519.28 | 2.14 |
| ZI: ~ trt * size + (1 \| Year) + (1 \| Plot / plantid)<br>Cond: ~ trt + size + (1 \| Year) + (1 \| Year:trt) + (1 \| Plot / plantid) | 17 | 8521.52 | 4.38 |
| ZI: ~ trt + size + (1 \| Year) + (1 \| Plot / plantid)<br>Cond: ~ trt * size + (1 \| Year) + (1 \| Year:trt) + (1 \| Plot / plantid) | 17 | 8525.30 | 7.16 |

| | | | |
|---|---|---|---|
| ZI: ~ trt * size + (1 \| Year) + (1 \| Year:trt) + (1 \| Plot / plantid)<br>Cond: ~ size + (1 \| Year) + (1 \| Plot / plantid) | 15 | 8517.76 | 0.62 |

**Table S13:** Models tested in model of number of umbels per plant. ZI corresponds to the zero-inflation portion of the model, i.e., the probability of producing zero umbels. Cond corresponds to the conditional portion of the model, i.e., the number of umbels produced per individual conditioned on flowering. Coefficients: `size` is size of plant in year of observation, `trt` is a categorical treatment, `Year` is year of observation. Asterisk (`*`) denotes both additive terms and an interaction term, while a colon (`:`) denotes only an interaction term, while parenthesized terms are model random effects. *p* denotes the number of parameters estimated. Bolded model is the final model used to estimate kernels.

| Model component | Parameter type | Coefficient name | Estimate | Std. error |
|---|---|---|---:|---:|
| Zero-inflation | Fixed | `Intercept` | 7.602 | 0.633 |
| | | `size` | -2.315 | 0.091 |
| | | `trtdrought` | 0.092 | 0.638 |
| | | `trtirrigated` | 0.558 | 0.643 |
| | Random | `Year` | 0.743 | - |
| | | `Year:trt` | 0.328 | - |
| | | `Plot` | 0.819 | - |
| | | `plantid:Plot` | 1.023 | - |
| Conditional | Fixed | `Intercept` | -4.306 | 0.191 |
| | | `size` | 1.090 | 0.038 |
| | Random | `Year` | 0.238 | - |
| | | `Plot` | 0.196 | - |
| | | `plantid:Plot` | 0.000 | - |

**Table S14:** Model coefficients in final model of inflorescence production used in kernel. Zero-inflation estimates correspond to the zero-inflation portion of the model, i.e., the probability of producing zero inflorescences. Conditional estimates correspond to the conditional portion of the model, i.e., the number of inflorescences produced per individual conditioned on flowering. Here, `size` corresponds to a continuous variable and `trt` corresponds to categorical variables with control as the reference group (i.e., term gives shift in intercept or slope for a treatment relative to the control).

| Model structures | *p* | AIC | ΔAIC |
|---|---|---|---|
| ZI: `Year + (1 \| Plot / plantid)`<br>Cond: `Year + (1 \| Plot / plantid)` | 13 | 15991.41 | 87.99 |
| ZI: `Year + (1 \| Plot / plantid)`<br>Cond: `size + Year + (1 \| Plot / plantid)` | 14 | 15961.87 | 58.45 |
| ZI: `size + Year + (1 \| Plot / plantid)`<br>Cond: `Year + (1 \| Plot / plantid)` | 14 | 15978.43 | 75.00 |
| ZI: `size + Year + (1 \| Plot / plantid)`<br>Cond: `size + Year + (1 \| Plot / plantid)` | 15 | 15949.24 | 45.82 |
| ZI: `size + Year + (1 \| Plot / plantid)`<br>Cond: `trt + size + Year + (1 \| Plot / plantid)` | 17 | 15949.16 | 45.73 |
| ZI: `trt + size + Year + (1 \| Plot / plantid)`<br>Cond: `size + Year + (1 \| Plot / plantid)` | 17 | 15951.88 | 48.45 |
| ZI: `trt + size + Year + (1 \| Plot / plantid)`<br>Cond: `trt + size + Year + (1 \| Plot / plantid)` | 19 | 15951.80 | 48.48 |
| ZI: `size + Year + (1 \| Plot / plantid)`<br>Cond: `trt*size + Year + (1 \| Plot / plantid)` | 19 | 15944.60 | 41.18 |
| ZI: `trt*size + Year + (1 \| Plot / plantid)`<br>Cond: `size + Year + (1 \| Plot / plantid)` | 19 | 15955.72 | 51.89 |
| ZI: `trt*size + Year + (1 \| Plot / plantid)`<br>Cond: `trt*size + Year + (1 \| Plot / plantid)` | 23 | 15950.72 | 47.29 |
| ZI: `trt + size + Year + (1 \| Plot / plantid)`<br>Cond: `trt*size + Year + (1 \| Plot / plantid)` | 21 | 15947.25 | 43.82 |
| ZI: `size + Year + (1 \| Plot / plantid)`<br>Cond: `trt*size + trt*Year + (1 \| Plot / plantid)` | 25 | 15948.11 | 44.69 |
| ZI: `size + trt*Year + (1 \| Plot / plantid)`<br>Cond: `trt*size + Year + (1 \| Plot / plantid)` | 27 | 15948.04 | 44.61 |
| ZI: `size + trt*Year + (1 \| Plot / plantid)`<br>Cond: `trt*size + trt*Year + (1 \| Plot / plantid)` | 33 | 15951.46 | 48.04 |
| ZI: `size + Year + (1 \| Plot / plantid)`<br>Cond: `n.umb + trt*size + Year + (1 \| Plot / plantid)` | 20 | 15945.84 | 42.41 |
| ZI: `n.umb + size + Year + (1 \| Plot / plantid)`<br>Cond: `trt*size + Year + (1 \| Plot / plantid)` | 20 | 15932.36 | 28.94 |

| | | | |
|---|---|---|---|
| ZI: n.umb + size + Year + (1 \| Plot / plantid)<br>Cond: n.umb + trt * size + Year + (1 \| Plot / plantid) | 21 | 15933.36 | 30.13 |
| ZI: n.umb + size + Year + (1 \| Plot / plantid)<br>Cond: phen + trt * size + Year + (1 \| Plot / plantid) | 21 | 15912.20 | 8.77 |
| ZI: n.umb + size + Year + (1 \| Plot / plantid)<br>Cond: phen^2 + trt * size + Year + (1 \| Plot / plantid) | 22 | 15914.18 | 10.76 |
| ZI: n.umb + size + Year + (1 \| Plot / plantid)<br>Cond: trt * size + trt * phen + Year + (1 \| Plot / plantid) | 23 | 15914.62 | 11.20 |
| ZI: n.umb + size + Year + (1 \| Plot / plantid)<br>Cond: trt * size + trt * phen^2 + Year + (1 \| Plot / plantid) | 26 | 15916.80 | 13.38 |
| ZI: n.umb + size + Year + (1 \| Plot / plantid)<br>Cond: trt * size + phen * Year + (1 \| Plot / plantid) | 24 | 15916.37 | 12.94 |
| ZI: n.umb + size + Year + (1 \| Plot / plantid)<br>Cond: trt * size + phen^2 * Year + (1 \| Plot / plantid) | 28 | 15923.13 | 19.71 |
| **ZI: phen + n.umb + size + Year + (1 \| Plot / plantid)<br>Cond: phen + trt * size + Year + (1 \| Plot / plantid)** | **22** | **15904.69** | **1.27** |
| ZI: trt * phen + n.umb + size + Year + (1 \| Plot / plantid)<br>Cond: phen + trt * size + Year + (1 \| Plot / plantid) | 26 | 15910.66 | 7.24 |
| ZI: n.umb + size + phen * Year + (1 \| Plot / plantid)<br>Cond: phen + trt * size + Year + (1 \| Plot / plantid) | 25 | 15904.39 | 0.97 |
| ZI: phen^2 + n.umb + size + Year + (1 \| Plot / plantid)<br>Cond: phen + trt * size + Year + (1 \| Plot / plantid) | 23 | 15903.42 | 0.00 |
| ZI: trt * phen^2 + n.umb + size + Year + (1 \| Plot / plantid)<br>Cond: phen + trt * size + Year + (1 \| Plot / plantid) | 29 | 15908.22 | 4.80 |
| ZI: n.umb + size + phen^2 * Year + (1 \| Plot / plantid)<br>Cond: phen + trt * size + Year + (1 \| Plot / plantid) | 29 | 15910.88 | 7.45 |

**Table S15:** Models tested in model of number of inflorescences per plant. ZI corresponds to the zero-inflation portion of the model, i.e., the probability that an inflorescence does not survive to the end of the season to produce seed. Cond corresponds to the conditional portion of the model, i.e., the number of seeds produced by a surviving inflorescence. Coefficients: `size` is size of plant in year of observation, `trt` is a categorical treatment, `Year` is year of observation, `n.umb` is the number of umbels (inflorescences) produced by the plant, and `phen` is the mean-centered flower emergence date of the plant. Asterisk (`*`) denotes both additive terms and an interaction term, a colon (`:`) denotes only an interaction term, superscript/caret (`^2`) denotes both a linear and quadratic term for the variable in question, and parenthesized terms are model random effects. *p* denotes the number of parameters estimated. Bolded model is the final model used to estimate kernels.

| Model | Parameter type | Coefficient name | Estimate | Std. error |
|---|---|---|---|---|
| Zero-inflation | Fixed | `Intercept` | 0.600 | 0.438 |
| | | `size` | -0.507 | 0.098 |
| | | `n.umb` | 0.128 | 0.037 |
| | | `Year2022` | 0.635 | 0.264 |
| | | `Year2023` | 0.853 | 0.267 |
| | | `Year2024` | 1.361 | 0.263 |
| | | `phen` | -0.018 | 0.006 |
| | Random | `Plot` | 0.145 | - |
| | | `plantid:Plot` | 0.029 | - |
| Conditional | Fixed | `Intercept` | 2.073 | 0.365 |
| | | `size` | 0.402 | 0.088 |
| | | `trtdrought` | 0.269 | 0.446 |
| | | `trtirrigated` | 1.331 | 0.492 |
| | | `Year2022` | 0.680 | 0.110 |
| | | `Year2023` | 0.414 | 0.112 |
| | | `Year2024` | 0.167 | 0.114 |
| | | `phen` | -0.149 | 0.003 |
| | | `trtdrought:size` | -0.124 | 0.107 |
| | | `trtdrought:irrigated` | -0.326 | 0.120 |
| | Random | `Plot` | 0.095 | - |
| | | `plantid:Plot` | 0.135 | - |

**Table S16:** Model coefficients in final model of seed production used in kernel. Zero-inflation estimates correspond to the zero-inflation portion of the model, i.e., the probability of inflorescence mortality before producing seed. Conditional estimates correspond to the conditional portion of the model, i.e., the number of seeds produced per inflorescence conditioned on the inflorescence surviving to produce seed. Here, `n.umb` corresponds to a continuous variable, and `Year` corresponds to a categorical variable with 2021 as the reference level (i.e., term gives shift in intercept for a year relative to 2021).

| Model structure | *p* | AIC | ΔAIC |
|---|---|---|---|
| `~ 1` | 2 | 379.95 | 98.19 |
| `~ (1 | Plot)` | 3 | 333.56 | 51.81 |
| `~ (1 | Year) + (1 | Plot)` | 4 | 284.66 | 2.91 |
| **`~ trt + (1 | Year) + (1 | Plot)`** | **6** | **281.76** | **0.00** |
| `~ trt + (1 | trt:Year) + (1 | Plot)` | 7 | 282.17 | 0.41 |

**Table S17:** Models fit to select the best model of new recruit size. Coefficients: `trt` is a categorical treatment and `Year` is year of census. A colon (**:**) denotes an interaction term and parenthesized terms are model random effects. *p* denotes the number of parameters estimated. Bolded model is the final model used to estimate kernels.

| Effect type | Coefficient name | Estimate | Std. error |
|---|---|---|---|
| Fixed | `Intercept` | 2.321 | 0.109 |
| | `trtdrought` | 0.185 | 0.104 |
| | `trtirrigated` | -0.100 | 0.106 |
| Random | `Plot` | 0.194 | - |
| | `Year` | 0.120 | - |
| | `Residual` | 0.351 | - |

**Table S18:** Model coefficients in final recruit size model used in kernels. Here, `trt` is a categorical with control as the reference group. The `Residual` random effect is the Gaussian error term, i.e., within-individual level standard deviation after accounting for all other model sources of variation.

| Contrast | Result set | Estimated Δλ | Sum of LTRE contributions | Relative error |
|---|---|---|---|---|
| Drought vs. control | Main | 0.015328 | 0.015254 | -0.005 |
| Drought vs. control | Mirrored | 0.015328 | 0.015355 | 0.002 |
| Irrigation vs. control | Main | 0.005720 | 0.005869 | 0.026 |
| Irrigation vs. control | Mirrored | 0.005720 | 0.005629 | -0.016 |

**Table S19:** Accuracy of LTRE results. Estimated Δλ is the difference in λ estimated from kernels generated for each treatment on its respective mean flowering date. Sum of LTRE contributions is the sum of contributions of both treatment and phenology effects across all vital rates. Relative error is estimated as [(estimated Δλ - summed LTRE contributions) / estimated Δλ].
